# A Comparative Genome-wide Transcriptome Analysis of Glucocorticoid Responder and Non-Responder Primary Human Trabecular Meshwork Cells

**DOI:** 10.1101/2021.09.21.461315

**Authors:** Kandasamy Kathirvel, Ravinarayanan Haribalaganesh, Ramasamy Krishnadas, Veerappan Muthukkaruppan, Colin E. Willoughby, Devarajan Bharanidharan, Srinivasan Senthilkumari

**Affiliations:** Department of Ocular Pharmacology, Aravind Medical Research Foundation, Madurai, Tamilnadu, India; Glaucoma Clinic, Aravind Eye Hospital, Madurai, Tamilnadu, India; Advisor, Aravind Medical Research Foundation, Madurai, Tamilnadu, India; Genomic Medicine, Biomedical Sciences Research Institute, Ulster University, Northern Ireland, United Kingdom; Department of Bioinformatics, Aravind Medical Research Foundation, Madurai, Tamilnadu, India

**Keywords:** Glucocorticoid-induced ocular hypertension, perfusion cultured human anterior segment, trabecular meshwork cells, gene expression, RNA-seq and candidate genes

## Abstract

The genome-wide gene expression analysis of primary human trabecular meshwork (HTM) cells with known glucocorticoid (GC) responsiveness was not reported earlier. Therefore, the purpose of this study was to investigate genes and pathways involved in the GC responsiveness in human trabecular meshwork (HTM) cells using RNA sequencing. A perfusion cultured human anterior segment *ex vivo* model was utilized to identify the induction of GC-induced ocular hypertension in one eye of a paired eyes after dexamethasone treatment based on the maximum intraocular pressure response and in the contralateral eye, HTM cells were isolated to classify GC-responder and non-responder cells. Some previously reported and unique genes and their associated pathways were identified in HTM cells in response to dexamethasone treatment versus vehicle control and more significantly in GC-responder and non-responder cells. This study will open up the possibility of identifying suitable molecular targets which have the potential to treat GC-induced ocular hypertension/glaucoma.

## 1. INTRODUCTION

Glucocorticoid (GC) therapy is the mainstay in the management of systemic and ocular autoimmune and inflammatory diseases. GC -induced ocular hypertension (GC-OHT) and glaucoma (GIG) are serious side-effects associated with long-term use of GC therapy. GC-OHT occurs in 40% of the population (GC-responders), out of which 6% are likely to develop glaucoma [1]. Individuals with GC responsiveness are at greater risk of developing primary open angle glaucoma (POAG) [2,3]. More than 90% of the patients with POAG exhibit GC responsiveness which can further aggravate the optic nerve damage leading to visual field loss [1]. The molecular mechanism responsible for the differential GC responsiveness is not clearly understood. Current glaucoma treatment therapy attempts to lower elevated intraocular pressure (IOP) with medications, laser or surgical treatment. However, currently there is no specific treatment option which specifically targets the pathogenic mechanisms responsible for the GC response in the trabecular meshwork (TM).

Several ‘omics’ studies attempted to identify the global changes in the expression of genes/proteins in TM cells in response to dexamethasone (DEX) treatment using cDNA or oligonucleotide arrays [4-11]. However, the overall findings were not consistent across these studies because of the difference in study methodology with different cell types, different duration and type of treatment and different microarray platforms used in these studies. Microarrays are limited due to the possibility of probe cross-hybridization, selection of specific probes and low detection thresholds. Moreover, the identification of novel transcripts and splice isoforms of the annotated genes are not possible due to its design with previously identified transcripts [12]. In addition, these studies did not include the details of IOP and GC responsiveness of the donor eyes from which TM cells were isolated. Therefore, it is difficult to detect if the changes in gene expression are directly or indirectly associated with GC-OHT pathogenesis or only reflect the effect of GCs on the TM. In a recent study, unique expression of genes and pathways were documented for GC-R and GC-NR bovine TM cells. In this study, perfusion cultured bovine anterior segment was utilized to identify eyes with induced OHT after DEX treatment in one eye of a paired eye and the primary bovine TM cell cultures were established in the contralateral paired eye with known GC responsiveness. The observed GC response rate for the bovine eyes in perfusion organ culture was found to be 36.8% [13]. However, the findings from bovine eyes may not be extrapolated to human GC responsiveness due to anatomical and physiological variations [14-16]. Therefore, in the present study the combination of both human organ-cultured anterior segment (HOCAS) *ex vivo* model and *in vitro* model of primary human TM (HTM) cells with known GC responsiveness were adopted to investigate the differential gene expression using genomewide transcriptome analysis with RNA-sequencing (RNA-seq).

Primary HTM cells with known GC responsiveness were assessed using the human organ cultured anterior segment (HOCAS) *ex vivo* model system to identify eyes with induced GC-OHT [13]. In a paired eye, one eye was used to assess the GC responsiveness following dexamethasone (DEX) treatment in HOCAS and the other eye was used to establish the primary HTM cells. This is the first study reporting a distinct gene signature with their associated pathways for GC-responder (GC-R) and non-responder (GC-NR) HTM cells. This study will open up the possibility of identifying suitable molecular targets which have the potential to treat GC-OHT/GIG.

## 2. RESULTS

### 2.1. Establishment of GC-R and GC-NR HTM cells

In this study, one eye of each set of paired eyes was used to assess the GC responsiveness in HOCAS after 100nM DEX treatment and the other eye was used to establish the primary HTM cells. Based on the IOP response, the HTM cells established from each donor eye was categorized as GC-R and GC-NR cells.

In HOCAS, DEX treatment caused an elevated IOP in 7/16 eyes (43.8%) [Mean Δ (±SEM) IOP: GC-R eyes: 13.8 ± 3.4 mmHg and GC-NR eyes: 0.91 ± 0.4 mmHg] (Fig. 1). The raw data of IOP of all the studied eyes are summarized in Supplementary Table 1.

**Fig.1.**
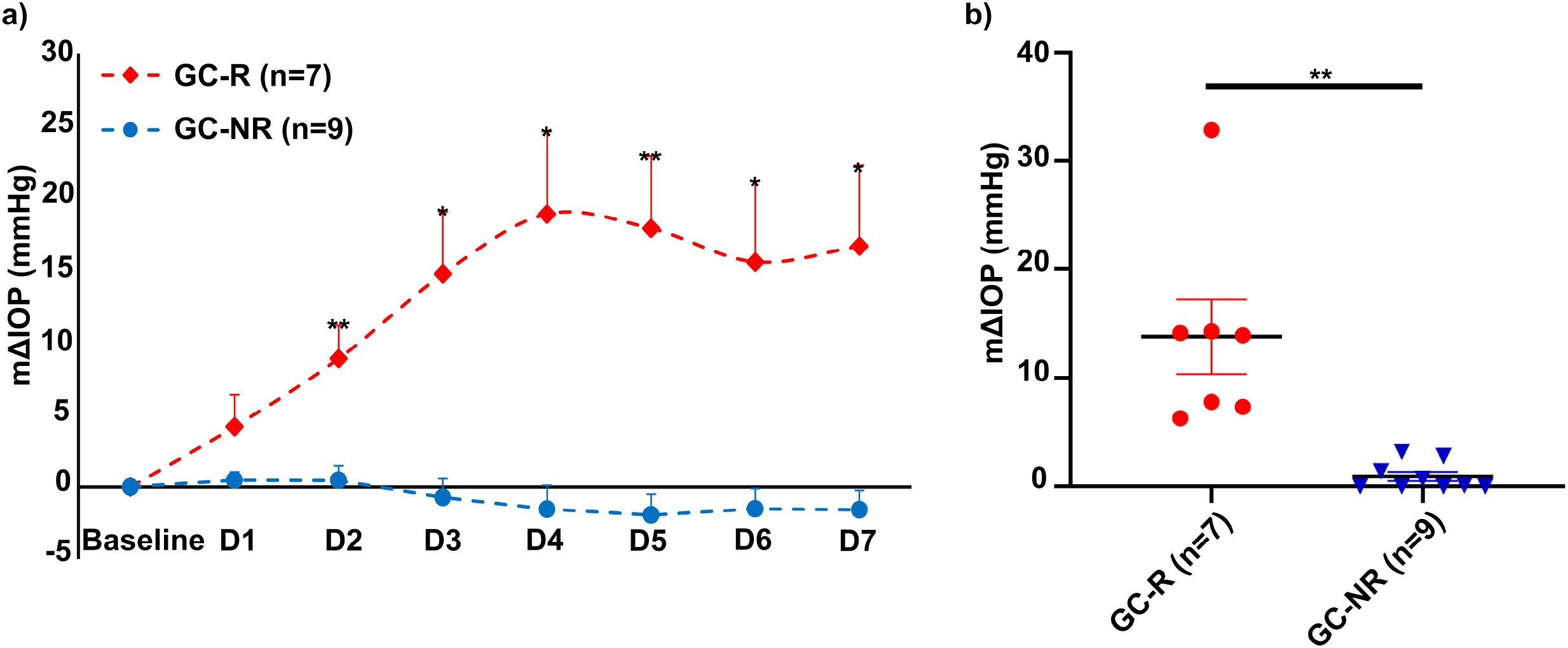
Effect of DEX on IOP. **a)** The mean ± SEM of ΔIOP of DEX-treated responder and non-responder eyes were plotted over time. The basal IOP on day 0 (before DEX treatment) was set at 0 mmHg. Treatment with DEX showed a significant elevated IOP in 7/16 eyes (Mean ± SEM - mΔIOP: 13.8 ± 3.41mmHg; Response rate: 43.8%). **b)** Frequency Plot of IOP data. The mΔIOP of GC-R and GC-NR groups were plotted. The mΔIOP was increased after DEX treatment in GC-R eyes as compared to GC-NR eyes. Data were analyzed by unpaired 2-tailed Student’s t test on each treatment day; \**p* <0.05; \*\**p* <0.001.

### 2.2. RNA Seq Data Quality

The fastQC evaluation of RNA seq revealed that the Phred score of all reads (forward and reverse) met the expected criteria of >30 (99.9% base call accuracy). After adapter and PCR duplicate trimming, approximately 5-7% of reads were excluded for further analysis. In addition to HTM cells with known GC responsiveness, primary HTM cells (n=2) cultured in DMEM media for 7 days was included to assess the effect of ETH on the expression of genes and found no significant changes between media treated and ETH treated cells (data not shown).

### 2.3. Differentially expressed genes of GC-R and GC-NR HTM cells

An average of 85.6% of mRNA reads were aligned with reference genome from all HTM cells used in the present study. The details of RNA-sequencing and alignment statistics are shown in Supplementary Table 2. The total number of genes identified in HTM cells of each donor eye ranged from 14,515 to 17,371. Principal component analysis of normalized data demonstrated that the DEX treated cells were dispersed from ETH treated cells (Supplementary Fig. 1). The expression of DEGs from GC-R (Group #1) and GC-NR (Group #2) HTM cells are represented in volcano plot (Fig. 2). In total, there were 616 and 216 DEGs in Group #1 (106 up-regulated; 510 down-regulated) and Group #2 (129 up-regulated; 87 down-regulated), respectively. There were 80 common genes found in Group #3 with absolute fold change (log2) value >2, and the *P value* < 0.05. Totally, 536 (56 up-regulated; 480 down-regulated) and 136 (78 up-regulated; 58 down-regulated) DEGs were found to be uniquely expressed only in GC-R (Group #4) and GC-NR (Group #5) HTM cells, respectively (Fig. 3). In Group #4, SAA4 (log FC=4.75), FRG2C (log FC=5.27) and NTRK2 (log FC=3.39) were significantly up-regulated and UPK3A (log FC=-8.48), RLN1 (logFC=-8.01) and NPY (logFC=-7.2) were significantly down-regulated whereas in Group #5, FAM107A (logFC=4.43), STEAP4 (logFC=4.57), RGCC (logFC=7.75) were significantly up-regulated and GRM5 (logFC= −5.05), SLC24A2 (logFC=-3.89) and GRIA2 (logFC=-3.82) were significantly down-regulated. In Group #3, the commonly expressed genes between Group #1 and #2 were SAA1, ZBTB16, FKBP5 and MYOC (up-regulated); and AQP1and LAMP3 (down-regulated). The top 50 DEGs from Group #1-Group #5 are shown in Supplementary Table 3a-3e, respectively.

**Figure 2.**
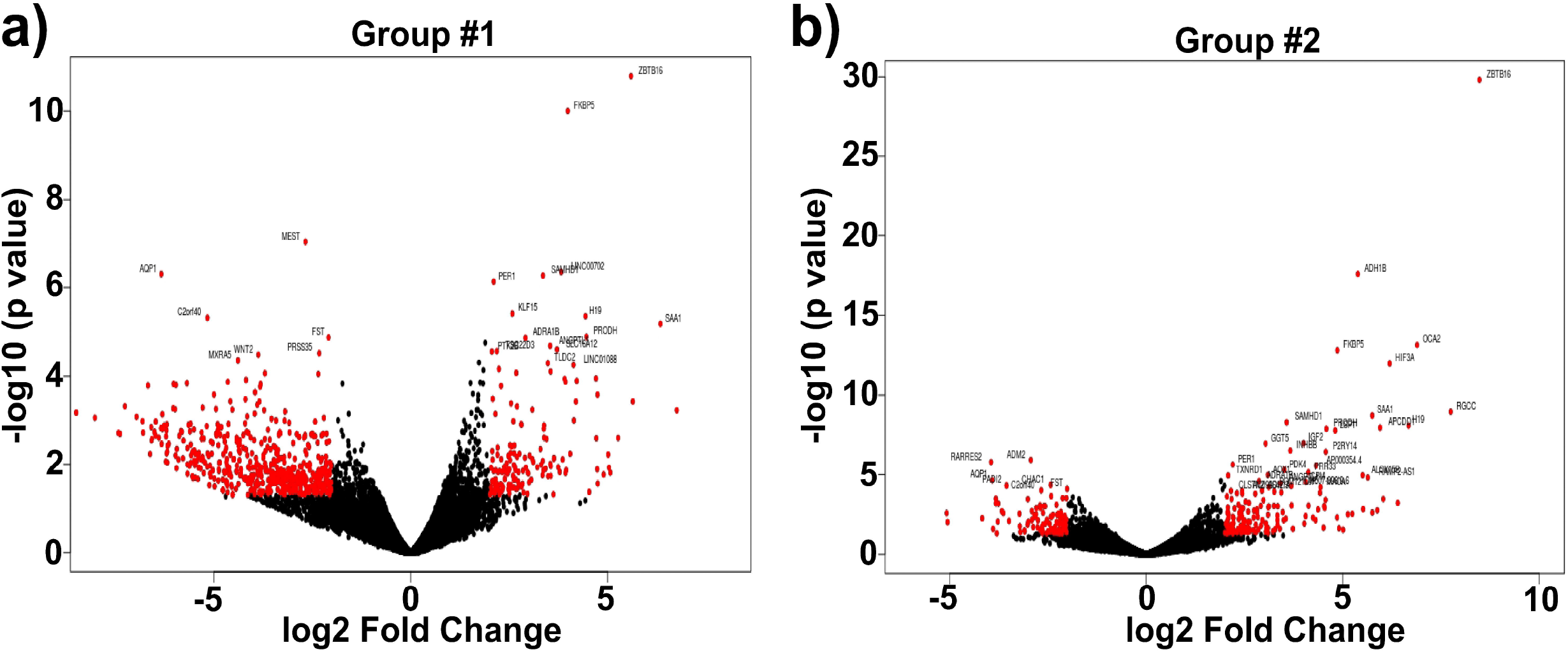
Volcano plot showing the Distribution of DEGs. The fold of change (log2) and p -value (-log10) of differentially expressed genes from (a) Group #1, (b) Group #2 are represented. Red color indicates the significantly dys-regulated genes with absolute fold change >2 and *p* -value: <0.05.

**Figure 3.**
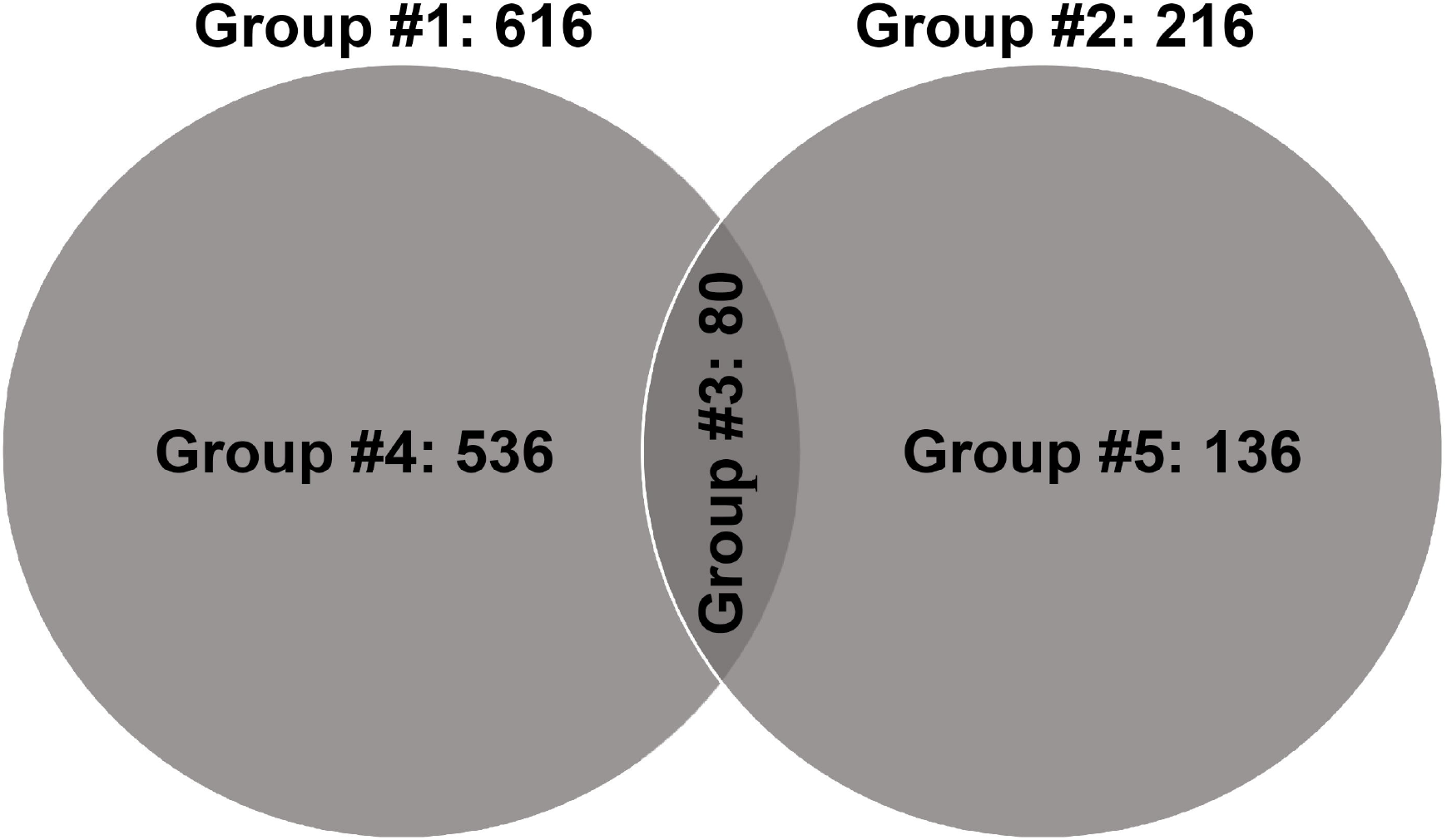
Venn diagram Showing Differentially Expression Groupings. DEG of three groups from RNA seq data are shown. Only genes with absolute fold change 2 and significant *p* value <0.05 were included in these groupings. Group #1: DEGs between ETH and DEX-treated cells of GC-R HTM cells, Group #2: DEGs between ETH and DEX-treated cells of GC-NR HTM cells, Group #3: Overlapping genes between Group #1 and Group #2; Group #4: uniquely expressed genes in GC-R and Group #5: uniquely expressed genes in GC-NR.

### 2.4. Pathway Analysis

Pathway analysis of DEGs from Group #3, Group #4 and Group #5 identified 75, 64 and 46 altered pathways, respectively. The altered pathways of each group were clustered into multiple functional categories. In Group #3, focal adhesion, WNT signaling, MAPK signaling, TGFβ signaling, drug metabolism cytochrome, cell adhesion, and pathways in cancer were found as most commonly enriched between GC-R and GC -NR HTM cells. Adherens junction, T cell receptor signaling, B cell receptor signaling, chemokine signaling pathway, regulation of actin cytoskeleton and TNF signaling pathways were enriched uniquely in GC-R HTM cells (Group #4). Interestingly, the predominant up-regulation of axon guidance, ECM-receptor interaction, focal adhesion, PI3K-Akt signaling pathways and down-regulation of WNT signaling, Calcium signaling pathway and vascular smooth muscle contraction, were identified in Group#4 than Group#5. The results of pathway analysis for the studied groups are summarized in Supplementary Table 4a - 4e.

### 2.5. Validation of DE genes by RT^2^-PCR Array

Out of 32 genes selected for PCR array, the expression pattern of 30 genes matched with RNA seq data, which further confirmed the reliability of these two techniques (Fig. 4a, 4b). Since, GAPDH and B2M showed significant changes to treatment in at-least one strain from each group, ACTB was used as a reference control.

**Fig.4a.**
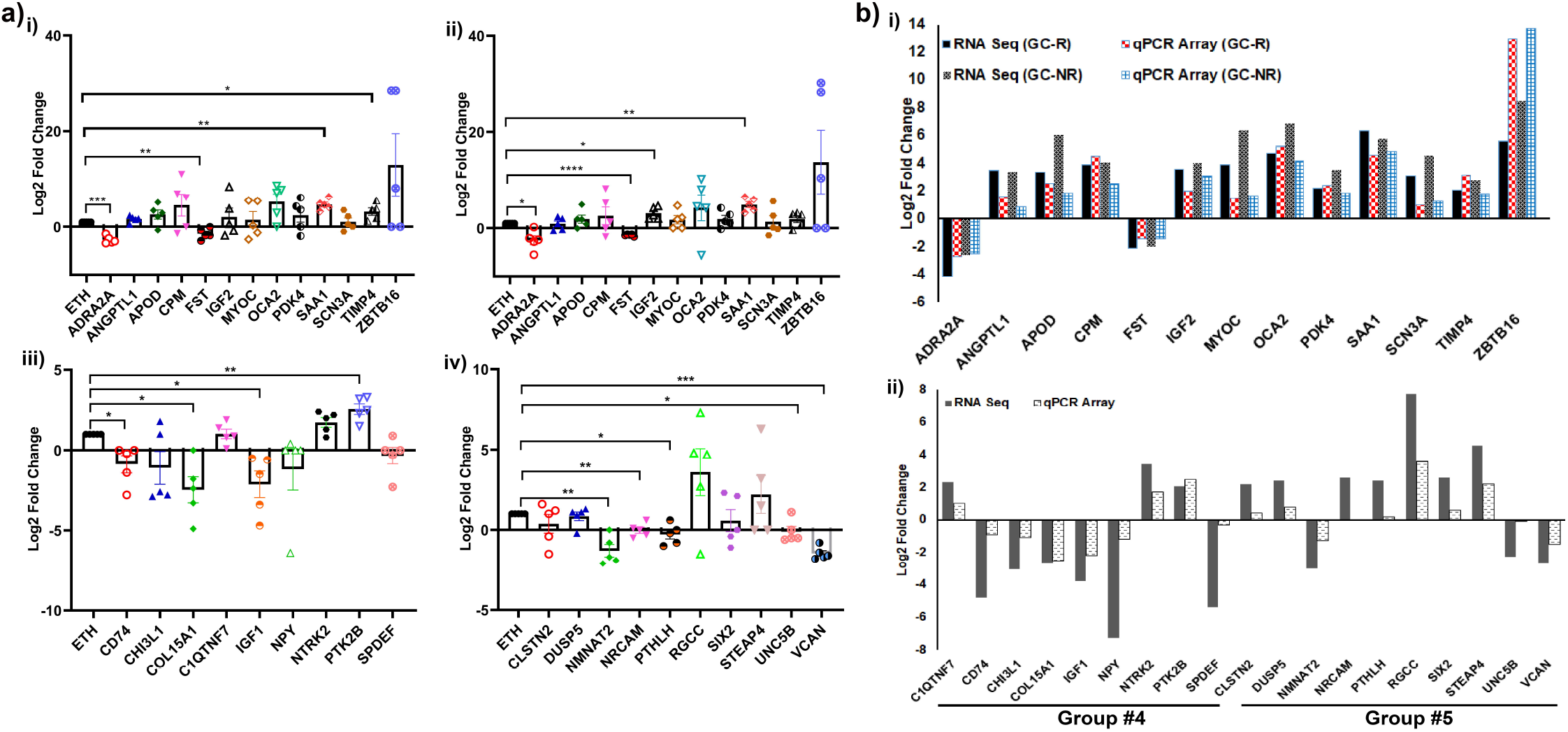
Validation of DEGs by RT^2^-PCR Array. Expression profile of selected genes identified from RNA-seq was validated by RT^2^-Profiler PCR array is shown. Primary HTM cells were treated with 100nM DEX or 0.1% ETH for 7 days. Total RNA was extracted, converted to cDNA and the expression profile of selected genes were carried out by RT^2^-PCR array as per the manufacturer’s instructions (Refer methods). Gene expressions were normalized to ACTB, and analyzed using the 2^-ΔΔCT^ method followed by Log_2_FC calculation. (i) Expression profile of selected genes from Group #3-GC-R; (ii) Group #3-GC-NR; (iii) Group #4 and (iv) Group #5 are shown. The data is represented as mean ± SEM. \**p* <0.05; \*\**p* <0.001; \*\*\**p* <0.0001. Paired 2-tailed Student’s t test. Fig. 4b. Validation of RNA sequencing Findings by RT^2^-PCR Array. Out of 32 genes, the expression pattern identified from RNA-seq of 30 genes matched with RT^2^-PCR array analysis. DEG Genes from (i) Group #3; (ii) Group #4 and group #5.

## 3. DISCUSSION

The pathophysiological mechanisms causing OHT/glaucoma associated with the use of GCs are not clearly understood. However, the dysfunction of TM associated with GC use as characterized by the structural and functional changes have been documented well [17]. Gene expression studies in cultured HTM cells [4-11] and perfusion cultured anterior segments using bovine eyes [13] after DEX treatment have identified differentially expressed genes associated with GC exposure in the TM (Supplementary Table 5). It is important to dissect generic alterations in gene expression associated with GC exposure in the TM from the genes involved in GC-induced OHT and GIG; that is, GC-responsiveness. Previous studies identified only a global change in the expression of genes in response to DEX treatment but not on the GC responsiveness related to humans [4-11]. The IOP response and the history of steroid sensitivity of the HTM cells/tissues derived from the donor eyes were not known in the earlier reported studies and hence there were inconsistencies in the expression of genes that are significantly altered. Therefore, in the present study the differentially expressed genes in primary HTM cells with known GC responsiveness was investigated using RNA-seq technology.

Uniquely in this study, the HOCAS *ex vivo* model was used to identify eyes with induced GC-OHT after DEX treatment based on the maximum IOP change (>5mmHg): GC responsiveness. Eyes could then be classified as GC-R or GC-NR based on a pathophysiological response in the HOCAS model with DEX treatment. Primary HTM cell cultures were established from the contralateral paired donor ryes after identification of GC responsiveness in HOCAS which provided cultured HTM cells with known GC responsiveness and represented a unique resource to explore the molecular basis of GC-OHT and GIG. A cell culture model was established as the yield of total RNA from individual TM tissue after HOCAS experiment was limited for RNA-seq experiment, hence the cultured HTM cells derived from GC-R and GC-NR eyes were further subjected for RNA-seq experiment after DEX treatment for 7 days. In order to validate the characteristics of primary HTM cells used in the present study, the expression of MYOC and CLAN formation after DEX induction were used. The cells with more than 50% MYOC positivity was used for all our analysis [18]. To our knowledge this is the first study reporting the differentially expressed genes in primary HTM cells with known GC responsiveness using RNA-seq technology.

The results of the present study revealed that an average of 16,022 genes were identified in cultured HTM cells and out of which the significantly altered genes in group #3, #4 and #5 were 80, 536 and 136, respectively. A higher number of significantly altered genes were found in GC-R cells (Group #1) (616 genes) as compared to GC-NR HTM cells (Group #2) (216 genes) in response to DEX treatment which indicates that both cells behaved differently to GC treatment.

A group of common genes which had been reported in previous studies to be up-regulated by DEX treatment in HTM cells were also found in our study (Supplementary Table 5). DEGs found in Group#4 are specifically relevant to GC-OHT/GIG as these genes were uniquely expressed DEGs of GC-R HTM cells. The apolipoprotein E (APOE) is the major lipoprotein in the central nervous system which plays an important role in the uptake and distribution of cholesterol within neuronal network. The polymorphism of APOE has been reported previously with the increased risk of POAG [19,20]. Our study showed that APOE was up-regulated only in GC-R cells (logFC= 1.86) and not in GC-NR group.

Neuropeptide Y (NPY) is involved in various physiological and homeostatic processes in both the central and peripheral nervous systems. It is expressed in the retina of both mammalian and non-mammalian species [21]. Immunohistochemical staining of NPY was also detected in drainage angle of mammalian eye [22]. NPY is reported to prevent neuronal cell death in retina induced by excitotoxic insults. Interestingly, in the present study it was highly down-regulated (logFC = −7.2) only in GC-R cells but not in GC-NR cells. Further studies are warranted to understand the role of NPY in dictating differential GC responsiveness in HTM cells.

Enrichment analysis with DEGs identified pathways previously reported in glucocorticoid treated TM cells and additional pathways enriched in the present study after DEX treatment (Supplementary Table 6a). Of 11 functional pathways that were reported to be significantly altered in human TM cells after exposure to DEX treatment versus a control medium, only 2 pathways such as cell adhesion and WNT signaling pathways were replicated in the present study [23]. The other reported pathways in earlier report were below the cut-off value and hence did not show any significant change in the present study [23]. Uniquely, in the present study we were able to sub-analyze the pathways enriched between the GC-R and GC-NR HTM cells (Supplementary Table 6b). One previous study had reported enriched pathways in bovine GC-R and GC-NR TM cells. The cell cycle and senescence pathways were highly significant between bovine GC-R and GC-NR TM cells. These pathways did not differ between GC-R and GC-NR HTM cells but interestingly our pathway analysis identified focal adhesion, WNT signaling, MAPK signaling, TGFβ signaling, drug metabolism cytochrome P450 and cell adhesion pathways to be associated with GC responsiveness (supplementary Table 6b). This variation may be attributed to the species variation and their corresponding responsiveness to DEX treatment. It is well documented in the perfusion cultured bovine eye model, the observed GC responsiveness was found to be 36.8% whereas in human eyes, it was between 30-40% as reported earlier by others and our group [13,24-26].

In the present study, it is interesting to note that WNT signaling was down-regulated in TM cells after DEX treatment as compared to vehicle control and upon comparing between TM cells of responder and non-responder cells, the WNT signaling genes (WNT2 (logFC = - 3.86), WNT4 (logFC = −2.48), WNT6 (logFC = −2.27), WNT7B (logFC = −3.84), WNT10A (logFC = −4.03), WNT10B (logFC = −3.03), WNT11 (logFC = −2.18)) and WNT signaling antagonist secreted frizzled-related proteins (sFRP2 (logFC = −4.09) and sFRP4 (logFC = −2.13)) were down-regulated in GC-R HTM cells (Group #4). In Group #5, WNT2 (logFC = −2.69) was identified as the only WNT signaling gene with down-regulated expression in GC-NR HTM cells. They differed in the intensity of gene expression. In general, abnormal WNT signaling has been associated with glaucoma and the expression of WNT signaling antagonist sFRP1 is reported to be up-regulated in glaucomatous TM cells [27]. The up-regulation of sFRP1 induced elevated IOP in both organ culture and murine models. The existence of active WNT signaling was documented in cultured HTM cells and the presence of which mediate ECM expression [28] and TM cell stiffening [29]. The WNT signaling small molecule inhibitor was effective in restoring the GC induced phenotypic changes in TM cells suggesting their potential applications in steroid induced glaucoma [30].

Several studies showed inhibitory crosstalk between GCs and transforming growth factor beta (TGF-β) in some but not all cell types [31,32], and inhibition of TGF-β signaling by GCs is known to be mediated either by reducing the bio-availability of TGF-β or by regulating the SMAD signaling [31-33]. Interactions between DEX and TGF-β signaling mediate GC-OHT as DEX activates TGF-β signaling inducing ER stress and ECM alterations resulting in IOP elevation [34]. The present study also identified the up-regulation of TGF-β signaling in both GC-R (logFC = 1.65) and GC-NR (logFC=1.38) human TM cells after DEX exposure which further confirms the observation of the previous study [34]. However, further investigations are warranted to decipher the functional role of TGF-β signaling in differential GC responsiveness.

The major strengths of this study include the use of TM cells derived from human cadaveric eyes with demonstrated GC responsiveness in HOCAS to investigate the transcriptome alterations. The observed GC response rate in perfusion cultured human cadaveric eyes of the present study was similar to previous study reported by other groups [25,35] and our group [24]. As age is known to be a risk factor for GC-OHT/glaucoma, the donor eyes from young age groups were excluded from the study. RNA-seq technology enabled us to identify unique genes and pathways and apply this experimental approach in GC-R and GC-NR human TM cells for the first time.

However, there are some limitations which need to be addressed. The TM tissues derived from human cadaveric eyes with known IOP response to GC treatment should have been ideal to investigate the transcriptome analysis related to GC responsiveness. Given the RNA quantity derived from TM tissues from donor eyes in our preliminary studies was not sufficient to run a robust RNA-seq analysis, cultured HTM cells derived from the contralateral paired eyes of known GC responsiveness were utilized in the present study. Usage of cultured cells might have contributed to variations in gene expression as compared to native tissues. In order to avoid such variability, HTM cells cultured in growth media for 7 days and gene expression profile was compared with HTM cells grown in medium containing 0.1% ethanol (vehicle control) and found no significant changes in gene expression pattern (data not shown). The history of glaucoma or any other ocular diseases of the human donor eyes used in the present study was not known and hence the expression profile of significantly altered genes between GC-R and GC-NR HTM cells might have been influenced. In addition, due to limited availability of the human donor eyes, some of the donor eyes used in the present study were above 48h, however, the viability of the tissues were maintained by storing them at 4°C immediately after enucleation until culture.

In conclusion, this is the first study reporting the differentially expressed genes in HTM cells with known GC responsiveness using RNA-seq technology. Utilizing perfusion cultured human cadaveric eyes in an *ex vivo* model enabled us to identify the induction of GC-OHT after DEX treatment and so classify HTM cells based on GC responsiveness: GC-R and GC-NR. Some previously reported and unique genes and their associated pathways were identified in TM cells in response to DEX treatment versus vehicle control; and more significantly in GC-R and GC-NR HTM cells. This study provides the potential to identify genes and proteins which are uniquely expressed by the GC responder eyes in Indian population. The further understanding of the molecular basis of HTM GC responsiveness could identify novel therapeutic approaches for GC-OHT and GIG which has direct clinical relevance given the use of glucocorticoids in ophthalmology and medicine in general.

## 4. Materials and Methods

### 4.1. Human Donor Eyes

Post-mortem human cadaveric eyes not suitable for corneal transplantation were obtained from the Rotary Aravind International Eye Bank, Aravind Eye Hospital, Madurai. The tissues were handled in accordance with the Declaration of Helsinki after getting approval from the standing Human Ethics Committee of the Institute. The donor eyes were enucleated within 5 h of death (mean elapsed time between death and enucleation was 2.75 ±1.58 h) and kept at 4° C in the moist chamber until culture. All eyes were examined under the dissecting microscope for any gross ocular pathological changes and only macroscopically normal eyes were used for the experiments.

In a set of paired eyes, one eye was used to establish HOCAS *ex vivo* model system to characterize GC responsiveness after DEX treatment and the other eye was used to establish primary HTM cultures from eyes with identified responsiveness [13] (Supplementary Fig. 2). The characteristics of donor eyes used for this study is summarized in Supplementary Table 7.

### 4.2. Human Organ Cultured Anterior Segment (HOCAS)

In a set of paired eyes, one eye was used to establish HOCAS by the method as described previously [13,24]. Briefly, after baseline equilibration (~72 h) one eye of each pair received 5 ml of 100 nM DEX for 7 days. The eye pressure was monitored continuously using pressure transducers (APT 300 Pressure Transducers, Harvard Apparatus, MA, USA) with data recorder (Power Lab system (AD Instruments, NSW, Australia) and LabChart Pro software (ver.8.1). The intraocular pressure (IOP) was calculated every hour as the average of 6 values recorded every 10 minutes, beginning 4 h before the drug infusion and continued for the duration of the culture. The average IOP in the 4 h before drug infusion was taken as baseline IOP for calculation. Mean IOP was calculated for every day after respective treatments. Then ΔIOP was calculated using the formula: (actual IOP averaged over 24 h - basal IOP of individual eyes on certain day)[13,36]. The increase in IOP in response to DEX treatment was examined for all treated eyes. The eyes were categorized as GC-responder (mean ΔIOP was > 5mmHg from the baseline) and non-responder eyes (mean ΔIOP< 5mmHg from the baseline) after DEX treatment for 7 days as described earlier [25].

### 4.3. Primary Human TM cell Strain with Known GC Responsiveness

The TM tissue was excised from the other eye of each set of paired eyes and the cell culture was established by extracellular matrix digestion method as described previously [37,38]. Primary HTM cells were grown at 37° C in 5% CO_2_ in low glucose Dulbecco’s modified Eagle medium (DMEM) with 15% fetal bovine serum, 5 ng/ml basic fibroblast growth factor and antibiotics. The primary HTM cells isolated from the other eye of each pair was characterized with aquaporin, myocilin and phalloidin staining by immunofluorescence analysis (Supplementary Fig. 3). HTM cell strain with more than 50% myocilin positivity were used for further experiments [18]. Confluent cultures of GC-R and GC-NR HTM cells were then treated with either 100nM DEX or 0.1% ethanol (ETH) as a vehicle control for 7 days and the medium was exchanged every other day. HTM cells from passages 2-4 were used for all experiments. At the end of the experiment, HTM cells from each GC-R (n=4) and GC-NR (n=4) were subjected to RNA extraction and RNA sequencing after DEX or 0.1% ETH treatment for 7 days.

### 4.4. RNA extraction and mRNA Sequencing

Total RNA was isolated from treated primary HTM cells by using the TRIZOL reagent (Sigma, MO, USA) as per the manufacturer’s instructions. RNA quantity and quality were assessed by NanoDrop 1000 spectrophotometer (Thermofisher Scientific, DE, UK), and TapeStation (Agilent Technologies, CA, USA), respectively. Additionally, the quality of RNA was observed by ratio of 28S and 18S ribosomal bands on 0.8 % agarose gel electrophoresis. The samples with RNA Integrity Number (RIN) value greater than 7 was used for RNA sequencing.

RNA sequencing for transcript profiling was performed at the Sandoor Lifesciences, Hyderabad, India. Briefly, 1μg of total RNA was used to enrich mRNA using NEB Magnetic mRNA isolation kit according to the manufacturer’s instructions (NEB, MA, USA). cDNA was synthesized and ligated to sequence adapters. The transcriptome library was prepared using NEB ultraII RNA library preparation kit as per the manufacturer’s recommended protocol (NEB, MA, USA) and the libraries then underwent size selection, PCR amplification and then PAGE purification. The final enriched libraries were purified and quantified by Qubit (Thermofisher Scientific, UK) and size analyzed by Bio-analyzer (Agilent Technologies, CA, USA). The resulting libraries were indexed and pooled then sequenced using Illumina Next Seq 500 (150 bp paired-end sequencing). Approximately 20-35 million reads were generated from each sample and obtained by de-multiplexing.

### 4.5. Mapping and Differential Expression Analysis

The quality of raw reads was assessed by FastQC toolkit (http://www.bioinformatics.babraham.ac.uk/projects/fastqc/) and adapter sequences were removed using bbduk.sh shell script from bbmap short read aligner (https://jgi.doe.gov/data-and-tools/bbtools/bb-tools-user-guide/bbmap-guide/). The pre-processed high-quality reads were then mapped with human reference genome assembly GRCh38/hg38 using HISAT2 by following default parameters [39]. The mRNA abundance in read counts were estimated using FeatureCounts [40]. mRNAs with less than 10 read counts were excluded from the further analysis. The read counts were then normalized using quantile strategy and the differential expression analysis with fragments per kilobase of exon per million (FPKM) values was performed by an R package: edgeR [41]. The mRNAs were considered as differentially expressed if the absolute fold change (log2) value was >2, and the P value < 0.05. For comparison, the Differentially expressed genes (DEGs) were segregated into five groups [13]: Group #1: DEGs between DEX and ETH treated GC-R HTM cells; Group #2 DEGs between DEX and ETH treated GC-NR HTM cells; and Group #3: DEGs that overlapping between Group #1 and Group #2; Group #4: Uniquely expressed DEGs of GC-R HTM cells (Group #1 minus Group #3); Group #5: Uniquely expressed DEGs of GC-NR HTM cells (Group #2 minus Group #3). We demonstrated our DEG analysis in Fig. 2a and b, for better understanding.

### 4.6. Pathway Enrichment Analysis

Pathways associated with DEGs were enriched using Database for annotation, visualization, and integrated discovery (DAVID) [42] with KEGG database. The pathways with fold enrichment of above 1 or below 1 with p value less than 0.05 were considered as significantly altered. The altered pathways were then clustered into their functional categories based on the molecular mechanisms.

### 4.7. Validation of RNA Seq by RT^2^-Profiler PCR Array

The expression of most upregulated and downregulated genes from Group #3, Group #4 and Group #5, identified in RNA-Seq was further validated by a RT^2^-Profiler PCR array (Qiagen, Hilden, Germany) as per the manufacturer’s instructions. List of genes taken for expression validation by RT^2^-Profiler PCR array is shown in Supplementary Table 8. Briefly, total RNA from TM cells after DEX treatment was isolated using RNAeasy mini kit (Qiagen, Hilden, Germany) and reverse transcribed into cDNA using RT2 first strand cDNA synthesis kit (Qiagen, Hilden, Germany), according to the manufacturer’s instructions. PCR Array was performed in a total volume of 25 μl containing 25 ng of total cDNA, 5x SYBR green master mix, loaded in each well containing gene specific probes along with reference controls: ACTB, B2M and GAPDH genes. The PCR array was performed by three steps of cycling program were 95°C for 10 min for 1 cycle, followed by 40 cycles of 95°C for 15s and 60°C for 60s, using the ABI-QuantStudio 5 (Applied Biosystems, MA, USA). The expression of genes in DEX treated HTM cells in logFC ratio was calculated by normalizing with reference control and vehicle control.

### 4.8. Statistical Analysis

Statistical analysis was carried out using Graph Pad Prism (ver.8.0.2) (Graph Pad software, CA, USA). All data are presented as mean ± SEM or otherwise specified. Statistical significance between two groups was analyzed using unpaired 2-tailed Student’s t test. *p* <0.05 or less was considered as statistically significant.

## Data Access

The raw mRNA sequencing data of HTM cells from each human donor eye used in the present study have been deposited publicly in NCBI-SRA under the BioProject PRJNA729873.

## Code Availability

The bioinformatics In-house pipeline used for mRNA sequencing data analysis in the present study have been submitted to GitHub in shell script (https://github.com/SenthilKumariLab/mRNA-seq-Analysis-Pipeline.git).

## Ethical approval

This study was approved by the standing Human Ethics Committee of Aravind Medical Research Foundation, Madurai, Tamilnadu, India (ID NO. RES2017006BAS).

## Consent for publication

Not applicable

## Author contributions

Conceptualization: SS; Methodology: SS, DB; Software: DB, KK; Validation: SS, DB; Formal Analysis: KK, RH, SS, DB; Investigation: KK, RH; Resources: SS, DB; Data Curation: SS, DB; Writing-Original Draft: KK, RH, SS, DB; Writing-Review and Editing: RK, VRM, CEW; Supervision: SS, DB; Project administration: SS; Funding acquisition: SS.

## Author Statement

All authors participated in the revision and approved the final version of the manuscript being submitted and declare no conflicting interest.

## Acknowledgements

The authors acknowledge the Rotary Aravind International Eye Bank, Aravind Eye Hospital, Madurai, India for providing human donor eyes for this study. we thank Dr. C. Gowri Priya, Scientist, Department of Immunology and Stem Cell biology, Aravind Medical Research Foundation. for rendering support for the acquisition of confocal image used in this study.

## References

[1] M.F. Armaly, Effect of Corticosteroids on Intraocular Pressure and Fluid Dynamics: I. The Effect of Dexamethasone* in the Normal Eye, Arch. Ophthalmol. 70 (1963) 482–491. https://doi.org/10.1001/archopht.1963.00960050484010.

[2] J.M. Lewis, T. Priddy, J. Judd, M.O. Gordon, M.A. Kass, A.E. Kolker, B. Becker, Intraocular pressure response to topical dexamethasone as a predictor for the development of primary open-angle glaucoma, Am. J. Ophthalmol. 106 (1988) 607–612. https://doi.org/10.1016/0002-9394(88)90595-8.

[3] J.D. Bartlett, T.W. Woolley, C.M. Adams, Identification of High Intraocular Pressure Responders to Topical Ophthalmic Corticosteroids, J. Ocul. Pharmacol. 9 (1993) 35–45. https://doi.org/10.1089/jop.1993.9.35.

[4] B.J. Fan, D.Y. Wang, C.C.Y. Tham, D.S.C. Lam, C.P. Pang, Gene expression profiles of human trabecular meshwork cells induced by triamcinolone and dexamethasone, Invest. Ophthalmol. Vis. Sci. 49 (2008) 1886–1897. https://doi.org/10.1167/iovs.07-0414.

[5] A. Nehmé, E.K. Lobenhofer, W.D. Stamer, J.L. Edelman, Glucocorticoids with different chemical structures but similar glucocorticoid receptor potency regulate subsets of common and unique genes in human trabecular meshwork cells, BMC Med. Genomics. 2 (2009) 58. https://doi.org/10.1186/1755-8794-2-58.

[6] A. Matsuda, Y. Asada, K. Takakuwa, J. Sugita, A. Murakami, N. Ebihara, DNA Methylation Analysis of Human Trabecular Meshwork Cells During Dexamethasone Stimulation, Invest. Ophthalmol. Vis. Sci. 56 (2015) 3801. https://doi.org/10.1167/iovs.14-16008.

[7] J.A. Faralli, H. Desikan, J. Peotter, et al., Genomic/proteomic analyses of dexamethasone-treated human trabecular meshwork cells reveal a role for GULP1 and ABR in phagocytosis, Mol. Vis. 25 (2019) 237–254.

[8] T. Ishibashi, Y. Takagi, K. Mori, et al., cDNA microarray analysis of gene expression changes induced by dexamethasone in cultured human trabecular meshwork cells., Invest. Ophthalmol. Vis. Sci. 43 (2002) 3691–7.

[9] W.R. Lo, L.L. Rowlette, M. Caballero, P. Yang, M.R. Hernandez, T. Borrás, Tissue Differential Microarray Analysis of Dexamethasone Induction Reveals Potential Mechanisms of Steroid Glaucoma, Invest. Ophthalmol. Vis. Sci. 44 (2003) 473. https://doi.org/10.1167/iovs.02-0444.

[10] Y.F. Leung, P.O.S. Tam, W.S. Lee, et al., The dual role of dexamethasone on antiinflammation and outflow resistance demonstrated in cultured human trabecular meshwork cells., Mol. Vis. 9 (2003) 425–39.

[11] F.W. Rozsa, D.M. Reed, K.M. Scott, et al., Gene expression profile of human trabecular meshwork cells in response to long-term dexamethasone exposure., Mol. Vis. 12 (2006) 125–41.

[12] J.D. Hoheisel, Microarray technology: beyond transcript profiling and genotype analysis, Nat. Rev. Genet. 7 (2006) 200–210. https://doi.org/10.1038/nrg1809.

[13] J.Y. Bermudez, H.C. Webber, B. Brown, T.A. Braun, A.F. Clark, W. Mao, A Comparison of Gene Expression Profiles between Glucocorticoid Responder and Non-Responder Bovine Trabecular Meshwork Cells Using RNA Sequencing, PLoS One. 12 (2017) e0169671. https://doi.org/10.1371/journal.pone.0169671.

[14] R.C. Tripathi, Ultrastructure of the exit pathway of the aqueous in lower mammals. (A preliminary report on the “angular aqueous plexus”), Exp. Eye Res. 12 (1971). https://doi.org/10.1016/0014-4835(71)90155-2.

[15] K. Erickson-Lamy, A.M. Schroeder, S. Bassett-Chu, D.L. Epstein, Absence of time-dependent facility increase (‘washout’) in the perfused enucleated human eye, Invest. Ophthalmol. Vis. Sci. 31 (1990) 2384–2388.

[16] P.A. Scott, D.R. Overby, T.F. Freddo, H. Gong, Comparative studies between species that do and do not exhibit the washout effect., Exp. Eye Res. 84 (2007) 435–43. https://doi.org/10.1016/j.exer.2006.10.015.

[17] M.E. Fini, S.G. Schwartz, X. Gao, et al., Steroid-induced ocular hypertension/glaucoma: Focus on pharmacogenomics and implications for precision medicine, Prog. Retin. Eye Res. 56 (2017) 58–83. https://doi.org/10.1016/j.preteyeres.2016.09.003.

[18] K.E. Keller, S.K. Bhattacharya, T. Borrás, et al., Consensus recommendations for trabecular meshwork cell isolation, characterization and culture, Exp. Eye Res. 171 (2018) 164–173. https://doi.org/10.1016/j.exer.2018.03.001.

[19] R. Liao, M. Ye, X. Xu, An updated meta-analysis: apolipoprotein E genotypes and risk of primary open-angle glaucoma., Mol. Vis. 20 (2014) 1025–36.

[20] N.M. Al-Dabbagh, N. Al-Dohayan, M. Arfin, M. Tariq, Apolipoprotein E polymorphisms and primary glaucoma in Saudis, Mol. Vis. 15 (2009) 912–919. http://www.molvis.org/molvis/v15/a95 (accessed May 22, 2021).

[21] A. Santos-Carvalho, F. Elvas, A.R. Álvaro, A.F. Ambrósio, C. Cavadas, Neuropeptide Y receptors activation protects rat retinal neural cells against necrotic and apoptotic cell death induced by glutamate, Cell Death Dis. 4 (2013) e636–e636. https://doi.org/10.1038/cddis.2013.160.

[22] T. Ohuchi, H. Tanihara, N. Yoshimura, S. Kuriyama, S. Ito, Y. Honda, Neuropeptide-induced [Ca2+]i transients in cultured bovine trabecular cells., Invest. Ophthalmol. Vis. Sci. 33 (1992) 1676–84.

[23] I. Liesenborghs, L.M.T. Eijssen, M. Kutmon, et al., The Molecular Processes in the Trabecular Meshwork After Exposure to Corticosteroids and in Corticosteroid-Induced Ocular Hypertension, Invest. Ophthalmol. Vis. Sci. 61 (2020) 24. https://doi.org/10.1167/iovs.61.4.24.

[24] R. Haribalaganesh, C. Gowri Priya, R. Sharmila, et al., Assessment of differential intraocular pressure response to dexamethasone treatment in perfusion cultured Indian cadaveric eyes., Sci. Rep. 11 (2021) 605. https://doi.org/10.1038/s41598-020-80112-8.

[25] A.F. Clark, K. Wilson, A.W. De Kater, R.R. Allingham, M.D. McCartney, Dexamethasone-induced ocular hypertension in perfusion-cultured human eyes, Invest. Ophthalmol. Vis. Sci. 36 (1995) 478–489. https://pubmed.ncbi.nlm.nih.gov/7843916/ (accessed July 9, 2020).

[26] M.F. Armaly, Statistical Attributes of the Steroid Hypertensive Response in the Clinically Normal Eye. I. The Demonstration of Three Levels of Response., Invest. Ophthalmol. Vis. Sci. 4 (1965) 187–97.

[27] W.-H. Wang, L.G. McNatt, I.-H. Pang, et al., Increased expression of the WNT antagonist sFRP-1 in glaucoma elevates intraocular pressure., J. Clin. Invest. 118 (2008) 1056–64. https://doi.org/10.1172/JCI33871.

[28] G. Villarreal, A. Chatterjee, S.S. Oh, D.-J. Oh, M.H. Kang, D.J. Rhee, Canonical Wnt Signaling Regulates Extracellular Matrix Expression in the Trabecular Meshwork, Invest. Ophthalmol. Vis. Sci. 55 (2014) 7433. https://doi.org/10.1167/iovs.13-12652.

[29] J.T. Morgan, V.K. Raghunathan, Y.-R. Chang, C.J. Murphy, P. Russell, Wnt inhibition induces persistent increases in intrinsic stiffness of human trabecular meshwork cells, Exp. Eye Res. 132 (2015) 174–178. https://doi.org/10.1016/j.exer.2015.01.025.

[30] S.D. Ahadome, C. Zhang, E. Tannous, J. Shen, J.J. Zheng, Small-molecule inhibition of Wnt signaling abrogates dexamethasone-induced phenotype of primary human trabecular meshwork cells.,Exp. Cell Res. 357 (2017) 116–123. https://doi.org/10.1016/j.yexcr.2017.05.009.

[31] U. Bolkenius, D. Hahn, A.M. Gressner, K. Breitkopf, S. Dooley, L. Wickert, Glucocorticoids decrease the bioavailability of TGF-β which leads to a reduced TGF-β signaling in hepatic stellate cells, Biochem. Biophys. Res. Commun. 325 (2004) 1264–1270. https://doi.org/10.1016/j.bbrc.2004.10.164.

[32] J.T. Schwartze, S. Becker, E. Sakkas, et al., Glucocorticoids Recruit Tgfbr3 and Smad1 to Shift Transforming Growth Factor-β Signaling from the Tgfbr1/Smad2/3 Axis to the Acvrl1/Smad1 Axis in Lung Fibroblasts, J. Biol. Chem. 289 (2014) 3262–3275. https://doi.org/10.1074/jbc.M113.541052.

[33] C.-Z. Song, X. Tian, T.D. Gelehrter, Glucocorticoid receptor inhibits transforming growth factor-beta signaling by directly targeting the transcriptional activation function of Smad3, Proc. Natl. Acad. Sci. 96 (1999) 11776–11781. https://doi.org/10.1073/pnas.96.21.11776.

[34] R.B. Kasetti, P. Maddineni, P.D. Patel, C. Searby, V.C. Sheffield, G.S. Zode, Transforming growth factor β2 (TGFβ2) signaling plays a key role in glucocorticoid-induced ocular hypertension, J. Biol. Chem. 293 (2018) 9854–9868. https://doi.org/10.1074/jbc.RA118.002540.

[35] M.F. Armaly, Effect of Corticosteroids on Intraocular Pressure and Fluid Dynamics: I. The Effect of Dexamethasone* in the Normal Eye, Arch. Ophthalmol. 70 (1963) 482–491. https://doi.org/10.1001/archopht.1963.00960050484010.

[36] W. Mao, T. Tovar-Vidales, T. Yorio, R.J. Wordinger, A.F. Clark, Perfusion-Cultured Bovine Anterior Segments as an Ex Vivo Model for Studying Glucocorticoid-Induced Ocular Hypertension and Glaucoma, Invest. Ophthalmol. Vis. Sci. 52 (2011) 8068–8075. https://doi.org/10.1167/IOVS.11-8133.

[37] W.D. Stamer, R.E.B. Seftor, S.K. Williams, H.A.M. Samaha, R.W. Snyder, Isolation and culture of human trabecular meshwork cells by extracellular matrix digestion, Curr. Eye Res. 14 (1995) 611–617. https://doi.org/10.3109/02713689508998409.

[38] S. Ashwinbalaji, S. Senthilkumari, C. Gowripriya, et al., SB772077B, A New Rho Kinase Inhibitor Enhances Aqueous Humour Outflow Facility in Human Eyes, Sci. Rep. 8 (2018). https://doi.org/10.1038/s41598-018-33932-8.

[39] D. Kim, J.M. Paggi, C. Park, C. Bennett, S.L. Salzberg, Graph-based genome alignment and genotyping with HISAT2 and HISAT-genotype, Nat. Biotechnol. 37 (2019) 907–915. https://doi.org/10.1038/s41587-019-0201-4.

[40] Y. Liao, G.K. Smyth, W. Shi, featureCounts: an efficient general purpose program for assigning sequence reads to genomic features, Bioinformatics. 30 (2014) 923–930. https://doi.org/10.1093/bioinformatics/btt656.

[41] M.D. Robinson, D.J. McCarthy, G.K. Smyth, edgeR: a Bioconductor package for differential expression analysis of digital gene expression data, Bioinformatics. 26 (2010) 139–140. https://doi.org/10.1093/bioinformatics/btp616.

[42] G. Dennis, B.T. Sherman, D.A. Hosack, et al., DAVID: Database for Annotation, Visualization, and Integrated Discovery, Genome Biol. 2003 49. 4 (2003) 1–11. https://doi.org/10.1186/GB-2003-4-9-R60.

