## Supplemental Figures for "A Comparative Genome-wide Transcriptome Analysis of Glucocorticoid Responder and Non-Responder Primary Human Trabecular Meshwork Cells"

### Slide 1
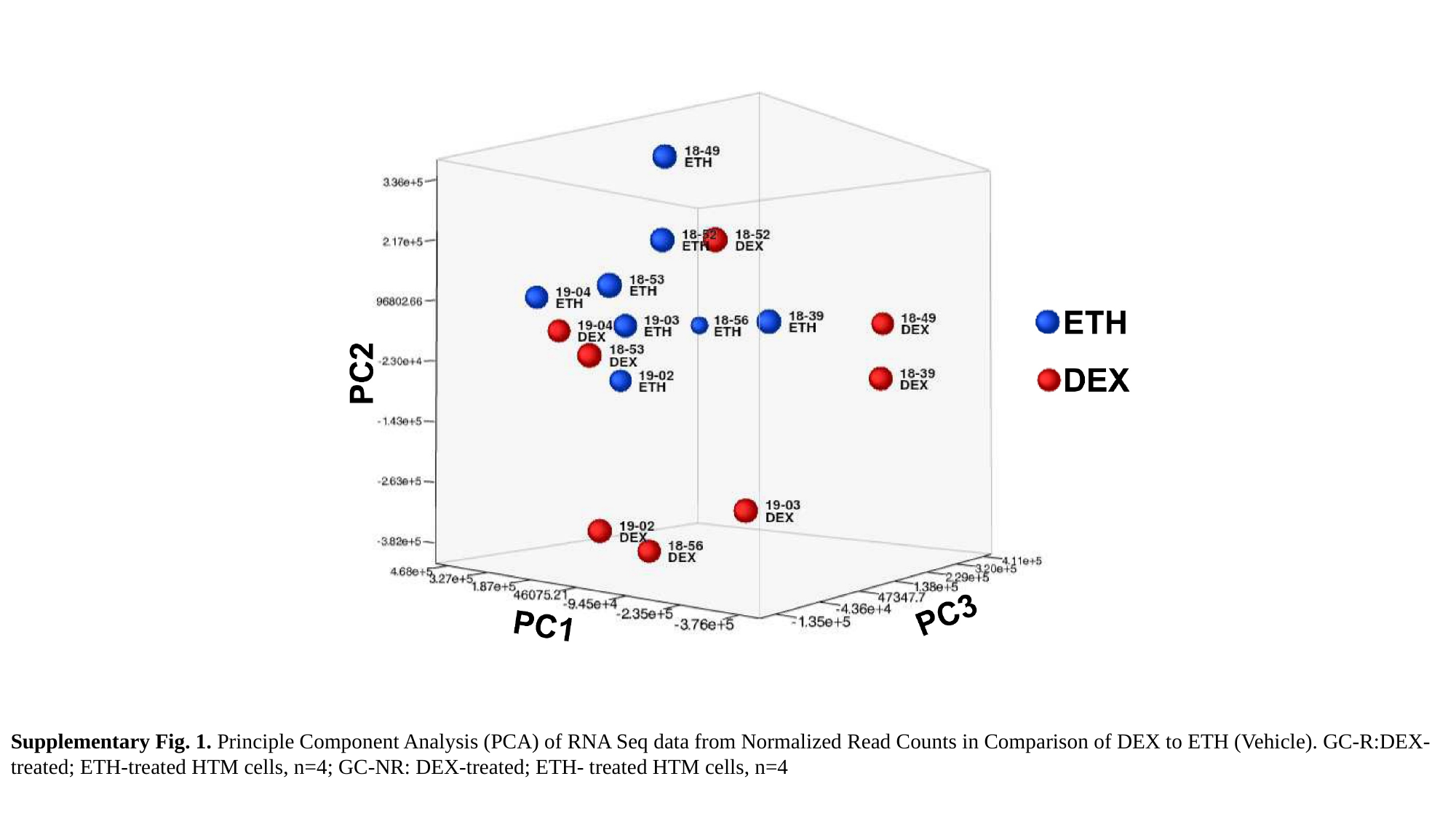

Supplementary Fig. 1. Principle Component Analysis (PCA) of RNA Seq data from Normalized Read Counts in Comparison of DEX to ETH (Vehicle). GC-R:DEX-treated; ETH-treated HTM cells, n=4; GC-NR: DEX-treated; ETH- treated HTM cells, n=4

### Slide 2
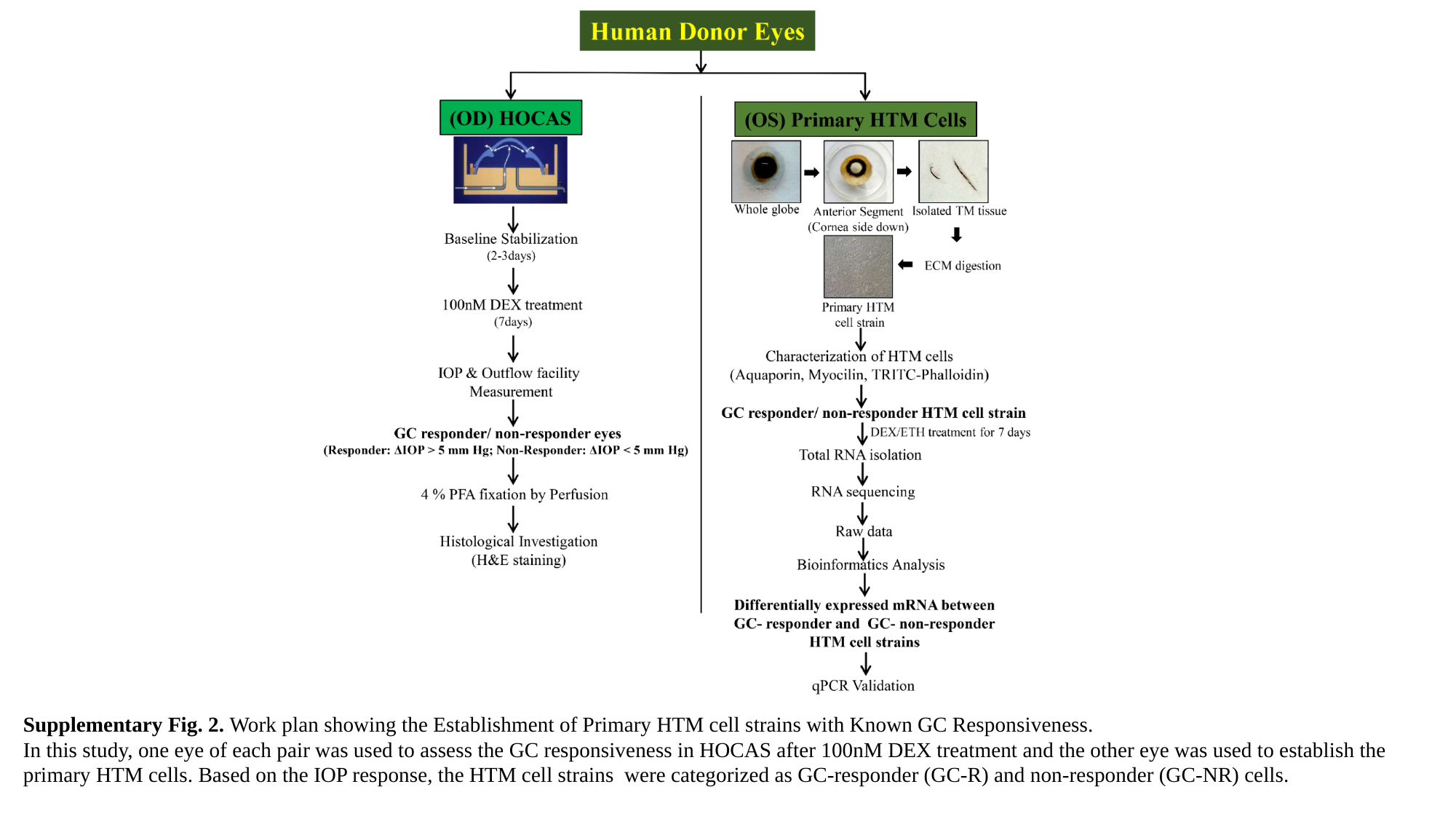

Supplementary Fig. 2. Work plan showing the Establishment of Primary HTM cell strains with Known GC Responsiveness.
In this study, one eye of each pair was used to assess the GC responsiveness in HOCAS after 100nM DEX treatment and the other eye was used to establish the primary HTM cells. Based on the IOP response, the HTM cell strains were categorized as GC-responder (GC-R) and non-responder (GC-NR) cells.

### Slide 3
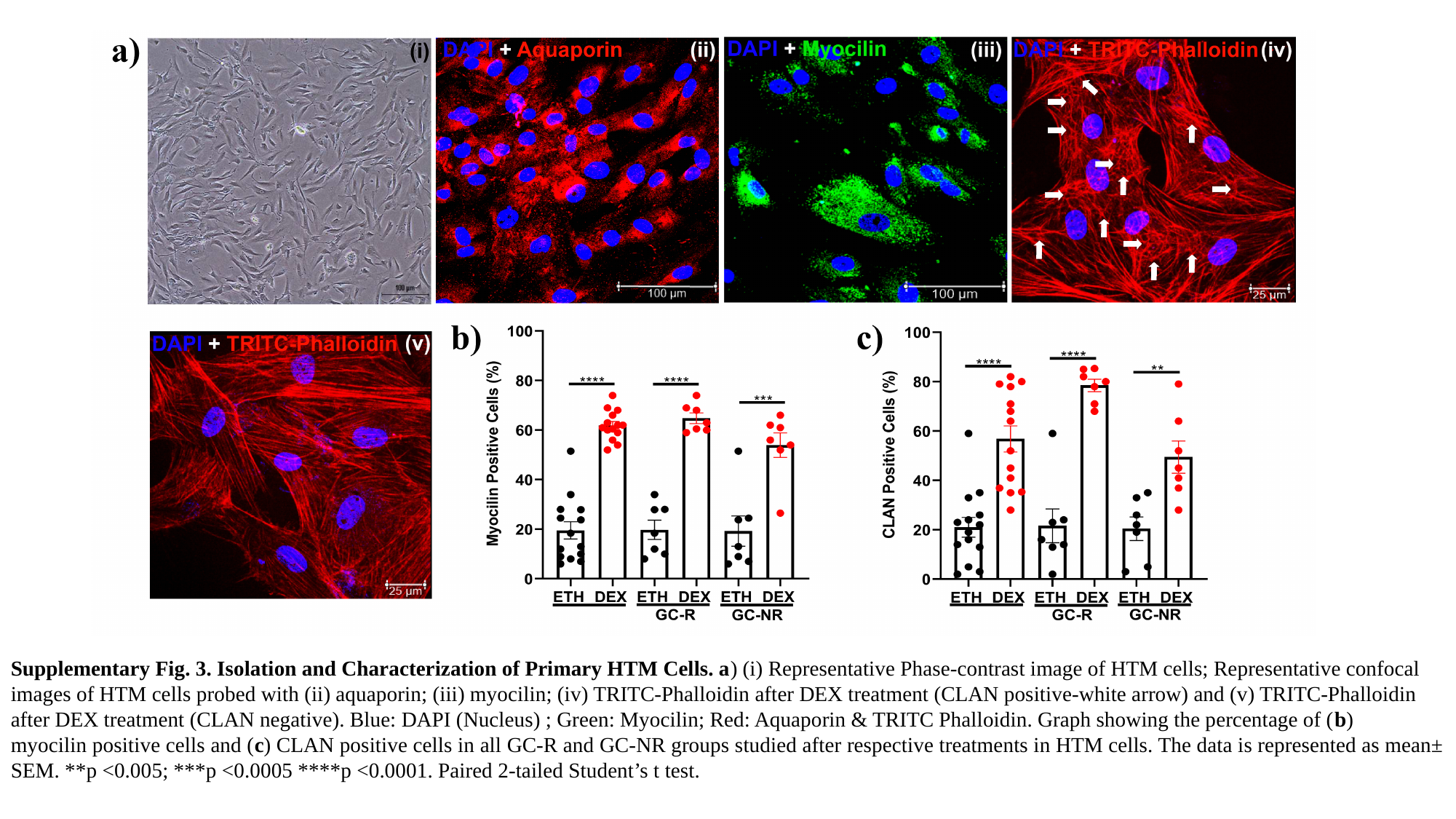

Supplementary Fig. 3. Isolation and Characterization of Primary HTM Cells. a) (i) Representative Phase-contrast image of HTM cells; Representative confocal
images of HTM cells probed with (ii) aquaporin; (iii) myocilin; (iv) TRITC-Phalloidin after DEX treatment (CLAN positive-white arrow) and (v) TRITC-Phalloidin
after DEX treatment (CLAN negative). Blue: DAPI (Nucleus) ; Green: Myocilin; Red: Aquaporin & TRITC Phalloidin. Graph showing the percentage of (b)
myocilin positive cells and (c) CLAN positive cells in all GC-R and GC-NR groups studied after respective treatments in HTM cells. The data is represented as mean± SEM. **p <0.005; ***p <0.0005 ****p <0.0001. Paired 2-tailed Student’s t test.
