## Supplemental Table for "A Comparative Genome-wide Transcriptome Analysis of Glucocorticoid Responder and Non-Responder Primary Human Trabecular Meshwork Cells"

**Supplementary Table 1:** The Raw Data of the IOP of Perfusion Cultured Indian Cadaveric Eyes

| **Dose group (nM)** | **R/NR** | **ID** | **Stabilization period (h)** | **Pressure change before and after drug treatment (mmHg)** | | | | | | | | **mΔ IOP (mmHg)** |
| --- | --- | --- | --- | --- | --- | --- | --- | --- | --- | --- | --- | --- |
|  |  |  |  | **D0** | **D1** | **D2** | **D3** | **D4** | **D5** | **D6** | **D7** |  |
| **100** | **R** | OCHD18-53 | 51.3 | 22.89 | 30.72 | 35.47 | 49.43 | 54.96 | 39.82 | 22.77 | 24.63 | 13.94 |
|  |  | OCHD18-57 | 40.8 | 17.57 | 16.47 | 17.94 | 18.62 | 18.16 | 22.73 | 35.99 | 37.21 | 6.30 |
|  |  | OCHD19-03 | 43.1 | 18.20 | 33.94 | 34.10 | 32.25 | 30.54 | 31.32 | 32.50 | 32.96 | 14.32 |
|  |  | OCHD19-04 | 35.5 | 21.62 | 21.71 | 23.49 | 28.47 | 33.03 | 33.80 | 33.61 | 31.81 | 7.80 |
|  |  | OCHD20-04 | 65.1 | 10.29 | 12.27 | 23.35 | 43.61 | 55.56 | 56.21 | 54.71 | 56.20 | 32.84 |
|  |  | OCHD20-09 | 37.8 | 11.94 | 15.47 | 24.34 | 20.79 | 22.40 | 23.70 | 12.74 | 15.63 | 7.36 |
|  |  | OCHD20-14 | 68.3 | 22.75 | 23.96 | 28.84 | 35.16 | 42.68 | 42.76 | 41.77 | 43.33 | 14.18 |
|  | Baseline IOP (IOP on day 0): mean= 17.89, SEM= 1.93, mΔ IOP: mean= 13.82, SEM= 3.4.10; N= 7 | | | | | | | | | | | |
|  | **NR** | OCHD18-29 | 62.3 | 22.61 | 23.48 | 24.43 | 23.22 | 22.20 | 23.13 | 23.05 | 23.82 | 0.72 |
|  |  | OCHD18-49 | 33.3 | 13.25 | 13.61 | 16.96 | 14.58 | 19.37 | 15.68 | 14.84 | 17.65 | 2.85 |
|  |  | OCHD18-52 | 48.1 | 20.75 | 20.86 | 23.22 | 24.63 | 25.66 | 25.12 | 25.32 | 22.98 | 3.22 |
|  |  | OCHD18-56 | 40.8 | 21.71 | 21.90 | 18.18 | 14.83 | 13.40 | 13.13 | 13.20 | 13.23 | 0 ^a^ |
|  |  | OCHD19-01 | 55.6 | 11.73 | 16.28 | 12.19 | 15.07 | 10.34 | 11.59 | 14.53 | 11.65 | 1.36 |
|  |  | OCHD19-02 | 27 | 12.50 | 13.18 | 16.60 | 13.34 | 12.42 | 11.78 | 10.09 | 10.61 | 0.07 |
|  |  | OCHD20-05 | 50.5 | 19.27 | 18.91 | 20.26 | 19.39 | 17.28 | 16.14 | 15.92 | 16.08 | 0 ^a^ |
|  |  | OCHD20-06 | 48.5 | 11.99 | 11.86 | 10.71 | 7.71 | 6.83 | 6.73 | 8.81 | 9.32 | 0 ^a^ |
|  |  | OCHD20-10 | 47.8 | 15.45 | 13.42 | 10.89 | 10.01 | 7.97 | 8.44 | 9.94 | 9.65 | 0 ^a^ |
|  | Baseline IOP (IOP on day 0): mean= 16.59, SEM= 1.7, mΔ IOP: mean= 0.91, SEM = 0.43; N= 9 | | | | | | | | | | | |

Note: ^a^ Minus values are given as 0; Day 0 represents the day before respective drug treatments; R - Responder; NR- Non-responder

**Supplementary Table 2:** Alignment Statistics of RNA Sequencing Data

| **S. No** | **R/NR** | **ID** | | **mRNA** | | | |
| --- | --- | --- | --- | --- | --- | --- | --- |
|  |  |  |  | **Total Reads**  **(In millions)** | **% Of Mapped Reads** | **% Of Unmapped Reads** | **No. of Genes** |
| 1 | **R** | 18-39 | ETH | 21.1 | 85.08 | 14.92 | 15765 |
|  |  |  | DEX | 29.3 | 84.79 | 15.11 | 17079 |
| 2 |  | 18-53 | ETH | 22.5 | 85.76 | 14.24 | 17371 |
|  |  |  | DEX | 36.2 | 82.11 | 17.89 | 16415 |
| 3 |  | 19-03 | ETH | 23.4 | 84.91 | 15.09 | 15353 |
|  |  |  | DEX | 30.5 | 87.00 | 13.00 | 16396 |
| 4 |  | 19-04 | ETH | 27.5 | 87.40 | 12.60 | 14515 |
|  |  |  | DEX | 30.7 | 85.38 | 14.62 | 16334 |
| 5 | **NR** | 18-49 | ETH | 26.8 | 84.30 | 15.70 | 15439 |
|  |  |  | DEX | 30.3 | 85.22 | 14.78 | 15637 |
| 6 |  | 18-52 | ETH | 23.1 | 84.42 | 15.58 | 14682 |
|  |  |  | DEX | 28.4 | 86.27 | 13.73 | 15997 |
| 7 |  | 18-56 | ETH | 24.8 | 86.28 | 13.72 | 15852 |
|  |  |  | DEX | 36.2 | 86.28 | 13.72 | 16654 |
| 8 |  | 19-02 | ETH | 23.4 | 88.60 | 11.40 | 16039 |
|  |  |  | DEX | 32.7 | 85.84 | 14.16 | 16828 |
|  |  |  | **Mean** | **28** | **86** | **14.4** | **16022** |

**Supplementary Table 3a**. List of top 50 Up/Down-regulated Genes from Group #1

| **Up-regulated genes** | | | |  | **Down-regulated genes** | | | |
| --- | --- | --- | --- | --- | --- | --- | --- | --- |
| **Gene** | **logFC** | **logCPM** | ***P* Value** |  | **Gene** | **logFC** | **logCPM** | ***P* Value** |
| SAA1 | 6.34 | 2.81 | 0.000 |  | UPK3A | -8.49 | 0.36 | 0.001 |
| ZBTB16 | 5.60 | 4.66 | 0.000 |  | RLN1 | -8.02 | 1.24 | 0.001 |
| FRG2C | 5.27 | -2.07 | 0.002 |  | FAM110D | -7.42 | -0.59 | 0.002 |
| NPSR1-AS1 | 5.07 | -2.19 | 0.016 |  | PRSS22 | -7.38 | -0.62 | 0.002 |
| BHLHE22 | 5.04 | 0.77 | 0.012 |  | NPY | -7.26 | 3.41 | 0.000 |
| SAA4 | 4.75 | -2.34 | 0.028 |  | GIMAP1 | -6.96 | -0.98 | 0.001 |
| SAA2 | 4.71 | 0.71 | 0.003 |  | ST14 | -6.81 | 1.52 | 0.001 |
| OCA2 | 4.71 | 1.61 | 0.000 |  | LRRC26 | -6.79 | 1.53 | 0.002 |
| PRODH | 4.46 | 3.17 | 0.000 |  | KRT15 | -6.67 | 3.39 | 0.000 |
| H19 | 4.44 | 7.60 | 0.000 |  | FXYD3 | -6.59 | 2.02 | 0.001 |
| PRR33 | 4.22 | 1.04 | 0.000 |  | PRAC1 | -6.58 | -0.09 | 0.003 |
| HIF3A | 4.20 | 4.11 | 0.000 |  | HOXB13 | -6.57 | 2.99 | 0.003 |
| P2RY14 | 4.14 | 1.13 | 0.001 |  | TPSB2 | -6.50 | 0.98 | 0.002 |
| LINC01088 | 4.14 | 1.46 | 0.000 |  | ACPP | -6.48 | 6.82 | 0.002 |
| FKBP5 | 3.99 | 7.23 | 0.000 |  | LCN2 | -6.47 | 0.44 | 0.001 |
| MYOC | 3.95 | 6.57 | 0.014 |  | KLK3 | -6.39 | 7.58 | 0.002 |
| CPM | 3.93 | 4.02 | 0.000 |  | KCNH2 | -6.34 | 0.38 | 0.002 |
| ANGPTL7 | 3.93 | 8.01 | 0.016 |  | AQP1 | -6.33 | 7.61 | 0.000 |
| LSP1 | 3.91 | 5.22 | 0.000 |  | DOC2B | -6.30 | -0.33 | 0.001 |
| LINC00702 | 3.82 | 3.76 | 0.000 |  | LINC01297 | -6.30 | 1.17 | 0.002 |
| TUSC5 | 3.79 | -1.37 | 0.015 |  | TPSAB1 | -6.26 | 0.71 | 0.001 |
| LEP | 3.75 | -0.16 | 0.012 |  | RBBP8NL | -6.22 | -1.54 | 0.009 |
| SLC16A12 | 3.72 | 1.63 | 0.000 |  | CHRNA2 | -6.21 | 1.31 | 0.002 |
| RNA5SP111 | 3.63 | -2.00 | 0.049 |  | SLC52A3 | -6.20 | -0.35 | 0.002 |
| IGF2 | 3.56 | 6.66 | 0.000 |  | KLK4 | -6.20 | 3.10 | 0.006 |
| KCNE1 | 3.55 | 0.74 | 0.010 |  | KRT5 | -6.19 | 2.48 | 0.001 |
| ANGPTL1 | 3.54 | 3.59 | 0.000 |  | MT1H | -6.18 | -1.57 | 0.005 |
| RPL7P57 | 3.49 | -2.06 | 0.019 |  | KLK2 | -6.17 | 6.41 | 0.004 |
| TLDC2 | 3.48 | 1.28 | 0.000 |  | CCDC64B | -6.03 | 0.46 | 0.001 |
| NTRK2 | 3.39 | 5.15 | 0.002 |  | SFN | -6.02 | 1.83 | 0.000 |
| ADH1B | 3.39 | 7.72 | 0.001 |  | DES | -5.98 | 6.03 | 0.001 |
| FMO2 | 3.38 | 3.38 | 0.002 |  | MSMB | -5.97 | 4.06 | 0.006 |
| SAMHD1 | 3.36 | 7.81 | 0.000 |  | COL9A1 | -5.96 | 2.75 | 0.000 |
| GIP | 3.36 | -1.29 | 0.008 |  | C2orf54 | -5.94 | -1.73 | 0.012 |
| APOD | 3.35 | 7.28 | 0.018 |  | PAGE4 | -5.94 | 0.47 | 0.004 |
| RN7SKP69 | 3.30 | -1.68 | 0.004 |  | EVX2 | -5.90 | -1.76 | 0.008 |
| MIR5690 | 3.22 | -1.75 | 0.012 |  | PLVAP | -5.87 | 1.64 | 0.002 |
| SCN3A | 3.09 | 0.86 | 0.008 |  | RAMP3 | -5.83 | 0.30 | 0.002 |
| TNFAIP8L3 | 3.09 | 3.20 | 0.001 |  | RAB25 | -5.81 | 0.28 | 0.002 |
| MAP1LC3C | 3.01 | 3.54 | 0.026 |  | HOXA9 | -5.78 | -0.78 | 0.007 |
| FHL5 | 2.99 | -0.36 | 0.016 |  | CHRM1 | -5.77 | 0.88 | 0.003 |
| SEMG1 | 2.97 | -1.56 | 0.047 |  | PDE9A | -5.68 | 1.33 | 0.000 |
| ABRA | 2.93 | 0.31 | 0.012 |  | SHH | -5.67 | -1.91 | 0.010 |
| ADRA1B | 2.91 | 2.49 | 0.000 |  | GIMAP7 | -5.67 | -0.28 | 0.004 |
| AFF2 | 2.90 | 1.40 | 0.007 |  | SPOCK3 | -5.64 | 1.74 | 0.001 |
| ADH1A | 2.85 | 1.95 | 0.014 |  | LCN6 | -5.61 | -1.94 | 0.011 |
| SLC16A10 | 2.84 | 1.28 | 0.028 |  | GNA15 | -5.60 | -0.92 | 0.002 |
| XRCC6P2 | 2.82 | -0.36 | 0.001 |  | HOXA10 | -5.59 | 0.11 | 0.001 |
| BCRP1 | 2.82 | -1.97 | 0.032 |  | MMP7 | -5.58 | 1.84 | 0.004 |
| HSPD1P11 | 2.81 | 1.79 | 0.001 |  | GATA5 | -5.58 | -0.87 | 0.009 |

Note: *Group #1*: DEGs between DEX and ETH treated GC-R HTM cells

**Supplementary Table 3b**. List of top 50 Up/Down-regulated Genes from Group #2

| **Up-regulated genes** | | | |  | **Down-regulated genes** | | | |
| --- | --- | --- | --- | --- | --- | --- | --- | --- |
| **Gene** | **logFC** | **logCPM** | ***P* Value** |  | **Gene** | **logFC** | **logCPM** | ***P* Value** |
| ZBTB16 | 8.48 | 4.59 | 0.000 |  | MXRA5Y | -5.08 | -0.47 | 0.003 |
| RGCC | 7.75 | 1.95 | 0.000 |  | GRM5 | -5.05 | -2.43 | 0.010 |
| OCA2 | 6.89 | 2.24 | 0.000 |  | RARRES2 | -3.94 | 3.36 | 0.000 |
| H19 | 6.67 | 8.47 | 0.000 |  | AQP1 | -3.91 | 4.05 | 0.000 |
| MYOC | 6.40 | 9.28 | 0.001 |  | SLC24A2 | -3.89 | -1.78 | 0.025 |
| HIF3A | 6.19 | 2.63 | 0.000 |  | LAMP3 | -3.82 | 0.03 | 0.000 |
| APOD | 6.03 | 8.49 | 0.000 |  | GRIA2 | -3.82 | 0.29 | 0.001 |
| APCDD1 | 5.95 | 2.15 | 0.000 |  | RBFOX1 | -3.80 | -1.82 | 0.048 |
| IGF2-AS | 5.88 | -1.83 | 0.002 |  | KRT17P1 | -3.77 | -1.24 | 0.009 |
| PNMT | 5.76 | -1.91 | 0.002 |  | TNFSF18 | -3.76 | 0.09 | 0.001 |
| SAA1 | 5.75 | 1.16 | 0.000 |  | GRID2 | -3.67 | -0.01 | 0.002 |
| RAMP2-AS1 | 5.64 | 0.83 | 0.000 |  | VCAN-AS1 | -3.63 | -1.95 | 0.003 |
| ALOX15B | 5.51 | -0.80 | 0.000 |  | PADI2 | -3.55 | 0.96 | 0.000 |
| ADH1B | 5.38 | 6.06 | 0.000 |  | PI16 | -3.49 | -0.82 | 0.008 |
| PTGDR2 | 5.24 | -2.22 | 0.003 |  | KLHDC7B | -3.22 | -0.85 | 0.003 |
| LINC00525 | 5.13 | -2.29 | 0.003 |  | LRRC15 | -3.21 | 4.55 | 0.017 |
| FKBP5 | 4.87 | 7.62 | 0.000 |  | IGFL2 | -3.20 | -2.15 | 0.040 |
| LSP1 | 4.81 | 4.97 | 0.000 |  | C1orf87 | -3.10 | -2.21 | 0.032 |
| PRODH | 4.58 | 2.59 | 0.000 |  | NPY6R | -3.09 | -2.22 | 0.036 |
| P2RY14 | 4.57 | 0.98 | 0.000 |  | NMNAT2 | -3.02 | 3.71 | 0.014 |
| STEAP4 | 4.57 | 3.40 | 0.000 |  | CLDN1 | -3.01 | 1.69 | 0.000 |
| SCN3A | 4.54 | 3.24 | 0.001 |  | ADM2 | -2.93 | 4.72 | 0.000 |
| SFTPC | 4.51 | -2.59 | 0.021 |  | RIMS3 | -2.91 | 2.92 | 0.002 |
| MAOA | 4.44 | 4.35 | 0.000 |  | CCND2-AS1 | -2.91 | -1.15 | 0.007 |
| FAM107A | 4.43 | 5.13 | 0.000 |  | IL32 | -2.91 | 2.34 | 0.007 |
| HMGN2P15 | 4.37 | -0.64 | 0.002 |  | TMEM63C | -2.84 | -1.45 | 0.003 |
| ANGPTL7 | 4.31 | 9.66 | 0.007 |  | COL19A1 | -2.79 | 0.34 | 0.001 |
| FHL5 | 4.24 | -0.81 | 0.005 |  | NGEF | -2.76 | 0.86 | 0.020 |
| SOAT2 | 4.13 | -0.86 | 0.001 |  | KAL1 | -2.75 | 2.99 | 0.007 |
| PRR33 | 4.13 | 0.77 | 0.000 |  | WNT2 | -2.70 | 0.87 | 0.001 |
| CPM | 4.06 | 3.13 | 0.000 |  | VCAN | -2.69 | 7.10 | 0.001 |
| SAA2 | 4.06 | -1.83 | 0.004 |  | CCND2-AS2 | -2.69 | -1.72 | 0.032 |
| TRAV39 | 4.03 | -0.91 | 0.012 |  | C2orf40 | -2.67 | 1.60 | 0.000 |
| IGF2 | 4.01 | 6.48 | 0.000 |  | NGFR | -2.65 | -1.76 | 0.020 |
| LGI3 | 3.70 | -0.07 | 0.001 |  | IPCEF1 | -2.64 | -2.20 | 0.047 |
| INHBB | 3.67 | 1.41 | 0.000 |  | VNN1 | -2.63 | -1.99 | 0.034 |
| SAMHD1 | 3.57 | 7.45 | 0.000 |  | ARHGAP9 | -2.62 | -0.48 | 0.003 |
| RPA4 | 3.56 | -2.16 | 0.006 |  | ADRA2A | -2.60 | 2.91 | 0.001 |
| PDK4 | 3.52 | 4.49 | 0.000 |  | RDH12 | -2.57 | -1.62 | 0.012 |
| ANKRD2 | 3.48 | 1.17 | 0.002 |  | PRSS35 | -2.57 | 3.69 | 0.001 |
| LINC00702 | 3.44 | 4.81 | 0.000 |  | MYHAS | -2.56 | -1.50 | 0.012 |
| ANGPTL1 | 3.42 | 2.28 | 0.000 |  | SLC14A1 | -2.53 | 0.08 | 0.047 |
| CHI3L2 | 3.34 | 3.45 | 0.015 |  | EPHA3 | -2.52 | 0.55 | 0.007 |
| SCARA5 | 3.34 | -0.26 | 0.004 |  | CNGA1 | -2.49 | -1.82 | 0.036 |
| GPRC5B | 3.26 | 5.02 | 0.000 |  | ELOVL2-AS1 | -2.46 | -1.70 | 0.012 |
| USP2 | 3.25 | 2.49 | 0.001 |  | KRT23 | -2.44 | -1.92 | 0.036 |
| TNNT3 | 3.12 | -0.91 | 0.001 |  | RGS7BP | -2.43 | -0.27 | 0.031 |
| POM121L9P | 3.12 | 2.50 | 0.000 |  | CHAC1 | -2.43 | 4.19 | 0.000 |
| ISM1 | 3.11 | -0.30 | 0.001 |  | ATP1A3 | -2.43 | -0.96 | 0.004 |
| NCAM1-AS1 | 3.10 | -2.39 | 0.046 |  | SEMA3D | -2.42 | 1.49 | 0.026 |

**Note:** *Group #2:* DEGs between DEX and ETH treated GC-NR HTM cells

**Supplementary Table 3c**. List of overlapping Genes from Group #3

| **Gene** | **Group#1** | | |  | **Group#2** | | |
| --- | --- | --- | --- | --- | --- | --- | --- |
|  | **logFC** | **logCPM** | ***P* Value** |  | **logFC** | **logCPM** | ***P* Value** |
| ZBTB16 | 5.60 | 4.66 | 0.000 |  | 8.48 | 4.59 | 0.000 |
| OCA2 | 4.71 | 1.61 | 0.000 |  | 6.89 | 2.24 | 0.000 |
| H19 | 4.44 | 7.60 | 0.000 |  | 6.67 | 8.47 | 0.000 |
| MYOC | 3.95 | 6.57 | 0.014 |  | 6.40 | 9.28 | 0.001 |
| HIF3A | 4.20 | 4.11 | 0.000 |  | 6.19 | 2.63 | 0.000 |
| APOD | 3.35 | 7.28 | 0.018 |  | 6.03 | 8.49 | 0.000 |
| SAA1 | 6.34 | 2.81 | 0.000 |  | 5.75 | 1.16 | 0.000 |
| ADH1B | 3.39 | 7.72 | 0.001 |  | 5.38 | 6.06 | 0.000 |
| FKBP5 | 3.99 | 7.23 | 0.000 |  | 4.87 | 7.62 | 0.000 |
| LSP1 | 3.91 | 5.22 | 0.000 |  | 4.81 | 4.97 | 0.000 |
| PRODH | 4.46 | 3.17 | 0.000 |  | 4.58 | 2.59 | 0.000 |
| P2RY14 | 4.14 | 1.13 | 0.001 |  | 4.57 | 0.98 | 0.000 |
| SCN3A | 3.09 | 0.86 | 0.008 |  | 4.54 | 3.24 | 0.001 |
| MAOA | 2.54 | 4.68 | 0.005 |  | 4.44 | 4.35 | 0.000 |
| ANGPTL7 | 3.93 | 8.01 | 0.016 |  | 4.31 | 9.66 | 0.007 |
| FHL5 | 2.99 | -0.36 | 0.016 |  | 4.24 | -0.81 | 0.005 |
| PRR33 | 4.22 | 1.04 | 0.000 |  | 4.13 | 0.77 | 0.000 |
| CPM | 3.93 | 4.02 | 0.000 |  | 4.06 | 3.13 | 0.000 |
| SAA2 | 4.71 | 0.71 | 0.003 |  | 4.06 | -1.83 | 0.004 |
| IGF2 | 3.56 | 6.66 | 0.000 |  | 4.01 | 6.48 | 0.000 |
| SAMHD1 | 3.36 | 7.81 | 0.000 |  | 3.57 | 7.45 | 0.000 |
| PDK4 | 2.24 | 5.55 | 0.000 |  | 3.52 | 4.49 | 0.000 |
| LINC00702 | 3.82 | 3.76 | 0.000 |  | 3.44 | 4.81 | 0.000 |
| ANGPTL1 | 3.54 | 3.59 | 0.000 |  | 3.42 | 2.28 | 0.000 |
| USP2 | 2.22 | 2.21 | 0.015 |  | 3.25 | 2.49 | 0.001 |
| MRO | 2.68 | 1.62 | 0.037 |  | 3.10 | 1.99 | 0.003 |
| AOX1 | 2.21 | 6.79 | 0.004 |  | 3.10 | 5.53 | 0.000 |
| GALNT15 | 2.26 | 4.60 | 0.014 |  | 3.04 | 4.52 | 0.012 |
| NEDD9 | 2.53 | 7.45 | 0.004 |  | 2.99 | 7.79 | 0.000 |
| ACA59 | 2.04 | -1.44 | 0.034 |  | 2.96 | -1.20 | 0.007 |
| ADH1A | 2.85 | 1.95 | 0.014 |  | 2.95 | 0.45 | 0.000 |
| FGFR4 | 2.50 | 3.10 | 0.029 |  | 2.92 | 1.76 | 0.001 |
| ADRA1B | 2.91 | 2.49 | 0.000 |  | 2.87 | 3.14 | 0.000 |
| PPP1R14A | 2.11 | 7.46 | 0.024 |  | 2.86 | 5.54 | 0.028 |
| XRCC6P2 | 2.82 | -0.36 | 0.001 |  | 2.84 | -0.33 | 0.005 |
| ABCA6 | 2.08 | 4.75 | 0.008 |  | 2.82 | 3.39 | 0.002 |
| TIMP4 | 2.07 | 1.11 | 0.025 |  | 2.78 | 2.41 | 0.000 |
| SLC16A12 | 3.72 | 1.63 | 0.000 |  | 2.75 | -0.57 | 0.002 |
| ADH4 | 2.54 | -1.36 | 0.029 |  | 2.74 | -0.95 | 0.005 |
| FAM46B | 2.15 | 4.36 | 0.001 |  | 2.53 | 4.96 | 0.001 |
| SLC16A10 | 2.84 | 1.28 | 0.028 |  | 2.47 | 0.64 | 0.011 |
| FPR1 | 2.70 | -0.01 | 0.007 |  | 2.46 | 0.79 | 0.024 |
| MOB3B | 2.61 | 2.14 | 0.001 |  | 2.45 | 2.31 | 0.020 |
| LINC01088 | 4.14 | 1.46 | 0.000 |  | 2.45 | 0.41 | 0.005 |
| FMO2 | 3.38 | 3.38 | 0.002 |  | 2.40 | 1.29 | 0.002 |
| TLDC2 | 3.48 | 1.28 | 0.000 |  | 2.38 | 1.05 | 0.003 |
| ANGPTL5 | 2.39 | 0.70 | 0.015 |  | 2.25 | 1.24 | 0.001 |
| OLAH | 2.57 | 2.30 | 0.004 |  | 2.20 | 0.37 | 0.004 |
| PER1 | 2.11 | 6.11 | 0.000 |  | 2.20 | 5.87 | 0.000 |
| TRPV6 | -3.05 | 1.87 | 0.019 |  | 2.17 | 0.67 | 0.014 |
| DKK2 | 2.46 | 3.59 | 0.016 |  | 2.17 | 3.54 | 0.017 |
| FST | -2.08 | 6.67 | 0.000 |  | -2.01 | 6.50 | 0.000 |
| TNFSF15 | -2.05 | 0.31 | 0.017 |  | -2.05 | -1.40 | 0.026 |
| CERS1 | -2.05 | -0.45 | 0.033 |  | -2.07 | -0.49 | 0.018 |
| ARSI | -2.42 | 3.36 | 0.001 |  | -2.13 | 3.10 | 0.000 |
| TNNT2 | -2.29 | 1.92 | 0.026 |  | -2.14 | -0.86 | 0.049 |
| BST2 | -4.64 | 1.92 | 0.000 |  | -2.20 | -0.98 | 0.012 |
| RGS16 | -2.34 | 5.10 | 0.007 |  | -2.21 | 5.59 | 0.028 |
| TAC3 | -3.33 | -1.20 | 0.002 |  | -2.23 | -1.29 | 0.033 |
| FNDC1 | -2.67 | 4.35 | 0.005 |  | -2.27 | 5.07 | 0.003 |
| SEMA6B | -2.30 | 1.97 | 0.016 |  | -2.28 | 1.15 | 0.001 |
| CPA4 | -2.42 | 3.57 | 0.002 |  | -2.34 | 2.68 | 0.034 |
| CILP2 | -2.02 | 0.07 | 0.026 |  | -2.34 | 0.13 | 0.024 |
| RGS7BP | -2.34 | -1.10 | 0.029 |  | -2.43 | -0.27 | 0.031 |
| KRT23 | -3.53 | 0.24 | 0.023 |  | -2.44 | -1.92 | 0.036 |
| SLC14A1 | -2.58 | 2.94 | 0.041 |  | -2.53 | 0.08 | 0.047 |
| PRSS35 | -2.32 | 3.30 | 0.000 |  | -2.57 | 3.69 | 0.001 |
| ADRA2A | -4.16 | 4.99 | 0.002 |  | -2.60 | 2.91 | 0.001 |
| NGFR | -2.06 | 0.83 | 0.041 |  | -2.65 | -1.76 | 0.020 |
| C2orf40 | -5.16 | 2.72 | 0.000 |  | -2.67 | 1.60 | 0.000 |
| WNT2 | -3.86 | 2.59 | 0.000 |  | -2.70 | 0.87 | 0.001 |
| NGEF | -2.86 | -0.57 | 0.016 |  | -2.76 | 0.86 | 0.020 |
| TMEM63C | -2.25 | -1.54 | 0.042 |  | -2.84 | -1.45 | 0.003 |
| LRRC15 | -2.42 | 2.19 | 0.026 |  | -3.21 | 4.55 | 0.017 |
| PI16 | -3.79 | 2.47 | 0.019 |  | -3.49 | -0.82 | 0.008 |
| TNFSF18 | -4.19 | 0.95 | 0.000 |  | -3.76 | 0.09 | 0.001 |
| LAMP3 | -3.62 | 0.92 | 0.002 |  | -3.82 | 0.03 | 0.000 |
| AQP1 | -6.33 | 7.61 | 0.000 |  | -3.91 | 4.05 | 0.000 |
| RARRES2 | -2.19 | 5.97 | 0.016 |  | -3.94 | 3.36 | 0.000 |
| MXRA5Y | -2.78 | -1.55 | 0.043 |  | -5.08 | -0.47 | 0.003 |

Note: *Group #3:* DEGs that overlapping between Group #1 and Group #2

**Supplementary Table 3d:** List of top 50 Up/Down-regulated Genes from Group #4

| **Up-regulated genes** | | | |  | **Down-regulated genes** | | | |
| --- | --- | --- | --- | --- | --- | --- | --- | --- |
| **Gene** | **logFC** | **logCPM** | ***P* Value** |  | **Gene** | **logFC** | **logCPM** | ***P* Value** |
| FRG2C | 5.27 | -2.07 | 0.002 |  | UPK3A | -8.49 | 0.36 | 0.001 |
| NPSR1-AS1 | 5.07 | -2.19 | 0.016 |  | RLN1 | -8.02 | 1.24 | 0.001 |
| BHLHE22 | 5.04 | 0.77 | 0.012 |  | FAM110D | -7.42 | -0.59 | 0.002 |
| SAA4 | 4.75 | -2.34 | 0.028 |  | PRSS22 | -7.38 | -0.62 | 0.002 |
| TUSC5 | 3.79 | -1.37 | 0.015 |  | NPY | -7.26 | 3.41 | 0.000 |
| LEP | 3.75 | -0.16 | 0.012 |  | GIMAP1 | -6.96 | -0.98 | 0.001 |
| RNA5SP111 | 3.63 | -2.00 | 0.049 |  | ST14 | -6.81 | 1.52 | 0.001 |
| KCNE1 | 3.55 | 0.74 | 0.010 |  | LRRC26 | -6.79 | 1.53 | 0.002 |
| RPL7P57 | 3.49 | -2.06 | 0.019 |  | KRT15 | -6.67 | 3.39 | 0.000 |
| NTRK2 | 3.39 | 5.15 | 0.002 |  | FXYD3 | -6.59 | 2.02 | 0.001 |
| GIP | 3.36 | -1.29 | 0.008 |  | PRAC1 | -6.58 | -0.09 | 0.003 |
| RN7SKP69 | 3.30 | -1.68 | 0.004 |  | HOXB13 | -6.57 | 2.99 | 0.003 |
| MIR5690 | 3.22 | -1.75 | 0.012 |  | TPSB2 | -6.50 | 0.98 | 0.002 |
| TNFAIP8L3 | 3.09 | 3.20 | 0.001 |  | ACPP | -6.48 | 6.82 | 0.002 |
| MAP1LC3C | 3.01 | 3.54 | 0.026 |  | LCN2 | -6.47 | 0.44 | 0.001 |
| SEMG1 | 2.97 | -1.56 | 0.047 |  | KLK3 | -6.39 | 7.58 | 0.002 |
| ABRA | 2.93 | 0.31 | 0.012 |  | KCNH2 | -6.34 | 0.38 | 0.002 |
| AFF2 | 2.90 | 1.40 | 0.007 |  | DOC2B | -6.30 | -0.33 | 0.001 |
| BCRP1 | 2.82 | -1.97 | 0.032 |  | LINC01297 | -6.30 | 1.17 | 0.002 |
| HSPD1P11 | 2.81 | 1.79 | 0.001 |  | TPSAB1 | -6.26 | 0.71 | 0.001 |
| C5AR2 | 2.76 | 0.11 | 0.006 |  | RBBP8NL | -6.22 | -1.54 | 0.009 |
| MARCH10 | 2.75 | -0.45 | 0.028 |  | CHRNA2 | -6.21 | 1.31 | 0.002 |
| KRT18P62 | 2.74 | -1.38 | 0.015 |  | SLC52A3 | -6.20 | -0.35 | 0.002 |
| HNRNPA3P11 | 2.73 | -1.59 | 0.025 |  | KLK4 | -6.20 | 3.10 | 0.006 |
| LINC00664 | 2.71 | -2.02 | 0.035 |  | KRT5 | -6.19 | 2.48 | 0.001 |
| KIAA1456 | 2.70 | 2.33 | 0.000 |  | MT1H | -6.18 | -1.57 | 0.005 |
| B3GALT2 | 2.68 | 3.10 | 0.000 |  | KLK2 | -6.17 | 6.41 | 0.004 |
| PLCE1-AS1 | 2.63 | 4.43 | 0.006 |  | CCDC64B | -6.03 | 0.46 | 0.001 |
| CYP51A1P2 | 2.62 | -1.04 | 0.034 |  | SFN | -6.02 | 1.83 | 0.000 |
| RAPGEF5 | 2.62 | 3.56 | 0.010 |  | DES | -5.98 | 6.03 | 0.001 |
| CCDC54 | 2.60 | -1.08 | 0.018 |  | MSMB | -5.97 | 4.06 | 0.006 |
| KLF15 | 2.58 | 4.75 | 0.000 |  | COL9A1 | -5.96 | 2.75 | 0.000 |
| ITGA10 | 2.55 | 4.93 | 0.000 |  | C2orf54 | -5.94 | -1.73 | 0.012 |
| PDLIM1P4 | 2.54 | -1.82 | 0.035 |  | PAGE4 | -5.94 | 0.47 | 0.004 |
| CYP7B1 | 2.53 | 1.20 | 0.008 |  | EVX2 | -5.90 | -1.76 | 0.008 |
| GPX3 | 2.51 | 7.49 | 0.013 |  | PLVAP | -5.87 | 1.64 | 0.002 |
| ANO3 | 2.48 | 0.01 | 0.006 |  | RAMP3 | -5.83 | 0.30 | 0.002 |
| NKAIN2 | 2.48 | 0.21 | 0.037 |  | RAB25 | -5.81 | 0.28 | 0.002 |
| LINC00547 | 2.47 | -0.51 | 0.013 |  | HOXA9 | -5.78 | -0.78 | 0.007 |
| ST7-AS2 | 2.44 | -1.06 | 0.009 |  | CHRM1 | -5.77 | 0.88 | 0.003 |
| FAM65C | 2.38 | 3.31 | 0.012 |  | PDE9A | -5.68 | 1.33 | 0.000 |
| ACTG1P3 | 2.33 | -1.48 | 0.038 |  | SHH | -5.67 | -1.91 | 0.010 |
| C1QTNF7 | 2.30 | 3.78 | 0.000 |  | GIMAP7 | -5.67 | -0.28 | 0.004 |
| GJA5 | 2.26 | 0.01 | 0.049 |  | SPOCK3 | -5.64 | 1.74 | 0.001 |
| FRMD3 | 2.21 | 1.81 | 0.006 |  | LCN6 | -5.61 | -1.94 | 0.011 |
| RN7SL608P | 2.19 | -1.55 | 0.028 |  | GNA15 | -5.60 | -0.92 | 0.002 |
| TSC22D3 | 2.19 | 8.12 | 0.000 |  | HOXA10 | -5.59 | 0.11 | 0.001 |
| SLC38A11 | 2.16 | 2.84 | 0.014 |  | MMP7 | -5.58 | 1.84 | 0.004 |
| LDHAL6B | 2.15 | -1.43 | 0.036 |  | GATA5 | -5.58 | -0.87 | 0.009 |
| FGF14 | 2.13 | 3.26 | 0.023 |  | LAD1 | -5.57 | -0.94 | 0.006 |

Note: *Group #4:* Uniquely expressed DEGs of GC-R HTM cells (Group #1 minus Group #3)

**Supplementary Table 3e:** List of top 50 Up/Down-regulated Genes from Group #5

| **Up-regulated genes** | | | |  | **Down-regulated genes** | | | |
| --- | --- | --- | --- | --- | --- | --- | --- | --- |
| **Gene** | **logFC** | **logCPM** | ***P* Value** |  | **Gene** | **logFC** | **logCPM** | ***P* Value** |
| RGCC | 7.75 | 1.95 | 0.000 |  | GRM5 | -5.05 | -2.43 | 0.010 |
| APCDD1 | 5.95 | 2.15 | 0.000 |  | SLC24A2 | -3.89 | -1.78 | 0.025 |
| IGF2-AS | 5.88 | -1.83 | 0.002 |  | GRIA2 | -3.82 | 0.29 | 0.001 |
| PNMT | 5.76 | -1.91 | 0.002 |  | RBFOX1 | -3.80 | -1.82 | 0.048 |
| RAMP2-AS1 | 5.64 | 0.83 | 0.000 |  | KRT17P1 | -3.77 | -1.24 | 0.009 |
| ALOX15B | 5.51 | -0.80 | 0.000 |  | GRID2 | -3.67 | -0.01 | 0.002 |
| PTGDR2 | 5.24 | -2.22 | 0.003 |  | VCAN-AS1 | -3.63 | -1.95 | 0.003 |
| LINC00525 | 5.13 | -2.29 | 0.003 |  | PADI2 | -3.55 | 0.96 | 0.000 |
| STEAP4 | 4.57 | 3.40 | 0.000 |  | KLHDC7B | -3.22 | -0.85 | 0.003 |
| SFTPC | 4.51 | -2.59 | 0.021 |  | IGFL2 | -3.20 | -2.15 | 0.040 |
| FAM107A | 4.43 | 5.13 | 0.000 |  | C1orf87 | -3.10 | -2.21 | 0.032 |
| HMGN2P15 | 4.37 | -0.64 | 0.002 |  | NPY6R | -3.09 | -2.22 | 0.036 |
| SOAT2 | 4.13 | -0.86 | 0.001 |  | NMNAT2 | -3.02 | 3.71 | 0.014 |
| TRAV39 | 4.03 | -0.91 | 0.012 |  | CLDN1 | -3.01 | 1.69 | 0.000 |
| LGI3 | 3.70 | -0.07 | 0.001 |  | ADM2 | -2.93 | 4.72 | 0.000 |
| INHBB | 3.67 | 1.41 | 0.000 |  | RIMS3 | -2.91 | 2.92 | 0.002 |
| RPA4 | 3.56 | -2.16 | 0.006 |  | CCND2-AS1 | -2.91 | -1.15 | 0.007 |
| ANKRD2 | 3.48 | 1.17 | 0.002 |  | IL32 | -2.91 | 2.34 | 0.007 |
| CHI3L2 | 3.34 | 3.45 | 0.015 |  | COL19A1 | -2.79 | 0.34 | 0.001 |
| SCARA5 | 3.34 | -0.26 | 0.004 |  | KAL1 | -2.75 | 2.99 | 0.007 |
| GPRC5B | 3.26 | 5.02 | 0.000 |  | VCAN | -2.69 | 7.10 | 0.001 |
| TNNT3 | 3.12 | -0.91 | 0.001 |  | CCND2-AS2 | -2.69 | -1.72 | 0.032 |
| POM121L9P | 3.12 | 2.50 | 0.000 |  | IPCEF1 | -2.64 | -2.20 | 0.047 |
| ISM1 | 3.11 | -0.30 | 0.001 |  | VNN1 | -2.63 | -1.99 | 0.034 |
| NCAM1-AS1 | 3.10 | -2.39 | 0.046 |  | ARHGAP9 | -2.62 | -0.48 | 0.003 |
| STOX1 | 3.10 | 2.24 | 0.023 |  | RDH12 | -2.57 | -1.62 | 0.012 |
| MYBPHL | 3.09 | -2.40 | 0.040 |  | MYHAS | -2.56 | -1.50 | 0.012 |
| C3 | 3.06 | 5.98 | 0.023 |  | EPHA3 | -2.52 | 0.55 | 0.007 |
| LINC00704 | 3.04 | -1.95 | 0.038 |  | CNGA1 | -2.49 | -1.82 | 0.036 |
| GGT5 | 3.04 | 6.46 | 0.000 |  | ELOVL2-AS1 | -2.46 | -1.70 | 0.012 |
| NR0B1 | 2.92 | -1.01 | 0.016 |  | CHAC1 | -2.43 | 4.19 | 0.000 |
| TRPC3 | 2.82 | 2.14 | 0.002 |  | ATP1A3 | -2.43 | -0.96 | 0.004 |
| FAM150B | 2.80 | -0.05 | 0.040 |  | SEMA3D | -2.42 | 1.49 | 0.026 |
| EDNRB | 2.79 | 1.64 | 0.014 |  | CTNNA3 | -2.37 | 1.35 | 0.003 |
| STAR | 2.76 | -1.13 | 0.002 |  | ERICH2 | -2.33 | -1.49 | 0.036 |
| WSCD1 | 2.74 | 0.98 | 0.034 |  | CCND2 | -2.32 | 2.26 | 0.019 |
| RPL23AP81 | 2.74 | -2.15 | 0.031 |  | LPHN3 | -2.30 | 2.21 | 0.043 |
| ALDH1L1-AS2 | 2.64 | -2.19 | 0.035 |  | INA | -2.30 | 2.04 | 0.038 |
| NRCAM | 2.61 | 3.46 | 0.001 |  | UNC5B | -2.28 | 5.69 | 0.009 |
| AKR1B15 | 2.60 | 0.70 | 0.009 |  | CA3 | -2.27 | -1.49 | 0.042 |
| B3GNT7 | 2.60 | 1.89 | 0.009 |  | RIMS2 | -2.25 | -1.14 | 0.045 |
| SYN2 | 2.60 | -0.79 | 0.002 |  | PLXNC1 | -2.23 | 2.68 | 0.003 |
| PLIN5 | 2.59 | -1.93 | 0.041 |  | LINC01133 | -2.21 | 2.67 | 0.003 |
| SIX2 | 2.58 | 4.07 | 0.019 |  | LRRN4CL | -2.19 | 2.88 | 0.005 |
| MTSS1 | 2.51 | 4.67 | 0.008 |  | KCNS1 | -2.17 | -0.42 | 0.010 |
| DGKG | 2.48 | 1.57 | 0.010 |  | KCTD16 | -2.17 | 2.93 | 0.002 |
| MREG | 2.47 | 2.69 | 0.001 |  | SLC7A5 | -2.14 | 6.66 | 0.004 |
| SPP1 | 2.45 | 0.63 | 0.000 |  | KCNMB2 | -2.12 | -1.25 | 0.033 |
| HEYL | 2.44 | 2.28 | 0.039 |  | BEX2 | -2.12 | -0.68 | 0.018 |
| TLE6 | 2.43 | 0.12 | 0.003 |  | CNTN6 | -2.11 | 1.58 | 0.010 |

Note: *Group #5:* Uniquely expressed DEGs of GC-NR HTM cells (Group #2 minus Group #3)

**Supplementary Table 4a.** List of Enriched Pathways in Group #1

| **Pathway** | **Fold Enrichment** | **P Value** | **Genes** |
| --- | --- | --- | --- |
| Phosphatidylinositol signaling system | 1.76 | 2E-03 | DGKG, PI4K2B, MTMR2, DGKE, DGKB, ITPR1, MTMR8, PIK3CD, ITPR2, PIK3CB, PIK3R1, PIK3C2A, MTMR6, MTMR7, MTM1, IMPA1, IMPA2, PPIP5K2, PIP4K2A, PLCE1, IP6K3, IPMK, INPP4B, PIKFYVE, PIK3CA, IMPAD1, PIK3C3, PLCB1, DGKI, DGKH |
| Focal adhesion | 1.73 | 7E-06 | ITGB1, GSK3B, FLT1, ITGB4, PIK3CD, PIK3CB, ARHGAP5, ACTB, IGF1R, PPP1CB, CCND3, AKT3, CAPN2, KDR, TNR, ITGAV, VAV3, PPP1R12A, ITGA4, ITGA2, ITGA1, ACTN4, PIK3CA, COL4A4, COL4A3, COL6A6, ITGA6, COL6A5, ITGA5, PPP1R12B, TLN1, MET, CRK, SOS2, BIRC2, SHC3, LAMA2, ROCK1, ROCK2, COL11A1, LAMA3, PDGFB, XIAP, PIK3R1, MYL12A, MYL12B, MAPK8, RAP1A, CHAD, SPP1, FLNA, MYL10, FLNB, PAK2, CAV2, CAV1, LAMB4, FN1, BRAF, PTK2, ITGA10, COL5A3 |
| TGF-beta signaling pathway | 1.65 | 1E-02 | TGIF1, SMAD2, SMAD4, BMPR2, ROCK1, ZFYVE9, NOG, CUL1, BMP8A, INHBB, INHBA, GDF6, LTBP1, SMAD5, ACVR2B, TGFBR1, TGFBR2, BMP2, ZFYVE16, RBL1, RPS6KB1, MYC, BMPR1B, NODAL |
| p53 signaling pathway | 1.63 | 3E-02 | GADD45B, APAF1, SERPINE1, CCNB3, CCND3, SESN3, CCNE2, RRM2B, ZMAT3, CHEK2, CASP3, CCNG2, CCNG1, CHEK1, MDM2, FAS, MDM4, ATM, ATR |
| mTOR signaling pathway | 1.59 | 7E-02 | PRKAA1, PRKAA2, CAB39, STRADA, IRS1, PIK3CD, BRAF, TSC1, PIK3CB, PIK3R1, PIK3CA, RPS6KB1, RPS6KA2, DDIT4, AKT3, RICTOR |
| Cell cycle | 1.58 | 6E-03 | CDKN1C, RB1, GSK3B, CCNH, PRKDC, CUL1, SMC3, CDC14B, CCNB3, CCND3, CHEK2, MYC, ORC3, RAD21, ORC2, CHEK1, CDC27, BUB3, BUB1, SMAD2, SMAD4, GADD45B, RBL2, CCNA1, STAG1, WEE1, RBL1, DBF4, STAG2, CCNE2, MDM2, ANAPC4, ATM, ATR |
| Adherens junction | 1.54 | 6E-02 | SMAD2, SMAD4, YES1, INSR, ACTN4, SORBS1, IQGAP1, TGFBR1, ACTB, TGFBR2, IGF1R, TJP1, SNAI1, SNAI2, CTNNA2, WASF1, MAP3K7, MET, WASF3 |
| Regulation of actin cytoskeleton | 1.51 | 1E-03 | ITGB1, CHRM2, NCKAP1, ITGB4, PIK3CD, PIK3CB, FGF1, FGF2, ACTB, FGF5, PPP1CB, CFL2, PIP4K2A, ITGAV, VAV3, PPP1R12A, ARHGEF12, ITGA4, ITGA2, ITGA1, ACTN4, ENAH, PIK3CA, ITGA6, ITGA5, PPP1R12B, CRK, SOS2, ARHGEF6, ROCK1, ROCK2, PDGFB, PIK3R1, IQGAP1, MYL12A, MYL12B, GNA13, NRAS, MYL10, WASF1, PAK2, FGF22, RDX, FN1, BRAF, ARPC5, SSH2, PTK2, PIKFYVE, DIAPH2, FGF14, APC, ITGA10, KRAS, FGFR4 |
| TNF signaling pathway | 1.51 | 2E-02 | ATF2, CEBPB, TNFAIP3, PIK3CD, PIK3CB, PIK3R1, CXCL2, MAPK8, BAG4, CASP3, AKT3, MAP3K8, DNM1L, MAP3K7, MAP3K5, EDN1, CHUK, CFLAR, FOS, TNFRSF1B, NFKBIA, ITCH, PIK3CA, TRAF5, FAS, TAB3, TAB2, BIRC2 |
| MAPK signaling pathway | 1.48 | 7E-04 | ATF2, ZAK, HSPB1, FGF1, FGF2, FGF5, MECOM, MYC, RPS6KA2, CASP3, AKT3, MAP3K8, MAP3K6, MAP3K7, MAP3K5, DUSP4, DUSP5, MEF2C, CHUK, DUSP1, CACNA2D3, CACNA2D4, FOS, TGFBR1, DUSP6, TGFBR2, CACNB2, PPM1A, PPM1B, TRAF6, RASA1, RASA2, RAPGEF2, CRK, SOS2, PDGFB, RASGRP2, RASGRP1, CACNG8, NRAS, PPP3R1, MAPK8, RAP1A, MAPK7, NTF3, FLNA, FLNB, CD14, PAK2, MAP4K3, FGF22, MAP3K2, NTRK2, GADD45B, NFATC3, BRAF, HSPA2, PPP5C, FGF14, TAOK1, NF1, FAS, KRAS, TAB2, FGFR4 |
| Pathways in cancer | 1.32 | 3E-03 | RB1, ITGB1, GSK3B, WNT2B, HHIP, PIK3CD, CBLB, PIK3CB, FGF1, ETS1, FGF2, IGF1R, FGF5, EDNRB, MECOM, MYC, CASP3, AKT3, ITGAV, JAK1, APPL1, ARHGEF12, CHUK, ITGA2, WNT5A, MITF, FOS, TGFBR1, TGFBR2, MSH6, PIK3CA, CCNE2, MSH2, MSH3, COL4A4, TRAF6, COL4A3, TRAF5, AGTR1, RARB, ITGA6, PLCB1, MET, CRK, SOS2, BIRC2, LAMA2, ROCK1, ROCK2, CUL2, LAMA3, PTGER3, PDGFB, GNAI3, LPAR1, WNT8B, XIAP, PIK3R1, CBL, RASGRP2, RASGRP1, HIF1A, FOXO1, GNA13, NRAS, MAPK8, TPR, CTNNA2, FGF22, RUNX1T1, EGLN1, SMAD2, STAT5A, SMAD4, ZBTB16, PTCH1, FZD6, LAMB4, FN1, BRAF, PTK2, NFKBIA, BMP2, FGF14, APC, GNAQ, GNB4, MDM2, FAS, KRAS |
| PI3K-Akt signaling pathway | 1.30 | 9E-03 | ITGB1, CHRM2, ATF2, GSK3B, FLT1, IRS1, ITGB4, PIK3CD, BRCA1, PIK3CB, FGF1, FGF2, IGF1R, GHR, FGF5, CCND3, MYC, AKT3, KDR, TNR, ITGAV, JAK2, IL6R, JAK1, ITGA4, CHUK, ITGA2, ITGA1, TSC1, PRLR, RBL2, PIK3CA, CCNE2, COL4A4, COL4A3, DDIT4, MTCP1, COL6A6, ITGA6, SGK3, COL6A5, ITGA5, MET, TLR4, SOS2, TLR2, PHLPP2, PRKAA1, PRKAA2, LAMA2, COL11A1, LAMA3, PDGFB, LPAR1, PIK3R1, FOXO3, NRAS, BCL2L11, PPP2R3C, CHAD, SPP1, FGF22, MCL1, ANGPT1, INSR, LAMB4, FN1, PPP2R3A, PTK2, FGF14, RPS6KB1, ITGA10, COL5A3, GNB4, MDM2, KRAS, PKN2, FGFR4 |
| Cell adhesion molecules (CAMs) | -1.94 | 5E-10 | CD86, CNTNAP2, CNTNAP1, ITGAM, ITGB2, ICAM2, F11R, ITGAL, CLDN1, PTPRF, ICAM1, SPN, CDH5, CDH3, CDH1, ITGB7, HLA-DOA, NEO1, CD34, ICOSLG, HLA-DPA1, HLA-B, HLA-C, HLA-A, HLA-F, CLDN11, CLDN6, CLDN5, CLDN4, CLDN3, CD8A, CLDN8, CLDN7, PECAM1, CDH15, HLA-DQB1, ITGA9, NLGN3, SELPLG, NLGN2, SDC4, NRXN1, SDC2, SDC3, NRXN2, GLG1, HLA-DMA, HLA-DMB, LRRC4C, HLA-DQA2, CD99, HLA-DQA1, LRRC4B, CD276, NTNG2, HLA-DRB5, VCAM1, CADM1, SELE, SELP, CD2, CD4, OCLN, PTPRC, SELL, CD6, HLA-DPB1, HLA-DRA, CNTN2, ESAM, SIGLEC1, CD22, HLA-DRB1 |
| Calcium signaling pathway | -1.68 | 2E-07 | RYR1, CHRM3, RYR2, MYLK2, CHRM1, CALML6, HTR2B, ATP2A3, ADRA1D, CALML3, ATP2A1, GRPR, ADRA1A, SLC8A2, MYLK4, PPP3CA, CYSLTR1, CYSLTR2, BDKRB2, CD38, BDKRB1, NOS1, PRKACA, PDGFRA, PRKCB, TNNC1, TACR2, AVPR1A, TACR1, ITPKB, ADCY9, LTB4R2, ADORA2B, STIM2, ORAI2, SLC25A5, PLCB2, SLC25A6, CAMK2B, PDE1B, PTGER1, PDE1A, CHRNA7, CAMK2A, PTAFR, CACNA1B, ADCY4, CACNA1A, CACNA1D, ADRB1, ADCY2, ADCY1, CACNA1E, ADCY7, CACNA1G, GNA14, HRH1, GNA15, PLN, PHKG1, ERBB3, TBXA2R, PHKG2, ERBB2, PLCG2, DRD1, NTSR1, NOS3, ATP2B3, GRIN2C, GRIN1, P2RX7, P2RX6, P2RX4, P2RX2, P2RX1, CAMK4, CALM3, PLCD3, PLCD4 |
| Axon guidance | -1.63 | 6E-05 | ROBO3, RND1, PPP3CA, RGS3, NCK2, RAC2, RAC3, PLXNC1, EPHB2, EPHB1, EPHB4, EPHB3, SEMA6B, EPHA7, SEMA6A, UNC5B, UNC5D, RHOD, EPHA1, NGEF, EPHA3, EPHA2, SEMA3D, SEMA3A, SEMA3B, SEMA3G, CXCR4, SEMA3E, SEMA3F, EFNA4, NTN3, EFNB1, PAK1, EFNB3, ABL1, PAK6, PLXNA1, SLIT3, SLIT2, SRGAP3, PAK3, LRRC4C, PAK4, NTNG2, SEMA4A, SEMA4D, NFATC2, SEMA4F, SEMA4G, EFNA1, EFNA3, CXCL12, CDK5, FES, PLXNB2 |
| Inflammatory mediator regulation of TRP channels | -1.50 | 5E-03 | CAMK2B, CALML6, CAMK2A, ADCY4, HTR2B, PIK3R3, CALML3, ADCY2, PIK3R2, ADCY1, ADCY7, PIK3CG, PIK3R5, ADCY5, HRH1, PLCG2, BDKRB2, BDKRB1, TRPM8, PRKACA, ASIC1, CYP2J2, NTRK1, PLA2G4F, PRKCH, PRKCB, PRKCE, PRKCD, PLA2G4C, IGF1, NGF, MAPK12, MAPK11, ADCY9, TRPV4, IL1B, F2RL1, CALM3, PLCB2 |
| ECM-receptor interaction | -1.47 | 1E-02 | LAMA5, TNXB, SDC4, LAMC3, LAMA1, ITGA2B, TNC, LAMC2, THBS2, THBS4, THBS3, VTN, RELN, SV2B, SV2A, DAG1, ITGB7, ITGB6, CD36, LAMB3, COL24A1, VWF, COL1A1, COL3A1, COL1A2, COL4A2, COL5A1, ITGA11, COL4A6, COL6A3, ITGA7, AGRN, CD44, ITGA9 |
| Wnt signaling pathway | -1.39 | 7E-03 | FZD10, PPP3CA, FRAT2, CCND2, SOX17, RAC2, RAC3, PRKACA, MMP7, PRKCB, CTNNBIP1, WNT9A, WNT16, DKK1, SFRP4, SFRP1, SFRP2, PLCB2, TP53, CAMK2B, CTBP1, TCF7, CAMK2A, LEF1, LRP5, CXXC4, NKD2, sFRP2, WNT11, WIF1, DVL1, TBL1Y, DVL2, DVL3, GPC4, WNT2, WNT3, WNT4, FZD1, WNT10B, WNT10A, FZD2, WNT3A, FZD7, WNT7B, FZD9, FZD8, NFATC2, NFATC1, VANGL2, BAMBI |
| cAMP signaling pathway | -1.33 | 5E-03 | RYR2, VIPR2, CHRM1, CALML6, PDE3B, CALML3, GLI1, PIK3CG, GLI3, CREB3L4, AKT2, CREB3L1, ADORA1, PDE4B, CNGA1, RAC2, RAC3, FFAR2, SOX9, PRKACA, PDE4C, NPY1R, SSTR1, ATP1B2, SSTR2, VAV1, VAV2, TIAM1, CREB3, ADCY9, RAPGEF3, CFTR, CREB5, CAMK2B, GRIA2, ADCYAP1R1, GIPR, NPR1, CAMK2A, ADCY4, PIK3R3, ATP1A3, CACNA1D, ADRB1, PIK3R2, ATP1A2, ADCY2, ADCY1, ADCY7, PIK3R5, ADCY5, PAK1, HCAR1, PLN, NPY, DRD1, GRIA4, BAD, BDNF, HTR1D, ATP2B3, NFATC1, GRIN2C, GRIN2B, TSHR, GRIN1, GRIN3B, CAMK4, FXYD2, CALM3 |
| Ras signaling pathway | -1.30 | 6E-03 | CALML6, FLT4, RASGRF2, RASGRF1, CALML3, PIK3CG, FGF7, SYNGAP1, FGF9, RASSF5, AKT2, RAC2, RAC3, PRKACA, PDGFRA, PLA2G4F, KSR1, PRKCB, HGF, PLA2G4C, NGF, PGF, PLA2G16, TIAM1, MRAS, KIT, RIN1, PLA1A, EPHA2, SHC2, SHC1, PLA2G1B, PDGFA, PIK3R3, PIK3R2, PLA2G5, RASAL3, EFNA4, RASGRP4, PIK3R5, PAK1, GNG4, GNG7, PDGFC, ABL1, PLCG2, PAK6, PAK3, RALGDS, PAK4, ANGPT4, NGFR, ANGPT2, EGF, BAD, PLA2G2A, VEGFB, VEGFC, IGF1, GRIN2B, GNG11, GRIN1, VEGFA, EFNA1, FGF17, EFNA3, FGF18, CALM3, GNB5, RGL1, FGF13, FGF12, FGFR3, RGL2, FGF11, FGFR2, FGF10, FGFR1 |

Note: Group #1: The pathways enriched from DEGs between ETH and DEX-treated GC- R HTM cells; Size: Number of genes enriched.

**Supplementary Table 4b.** List of Enriched Pathways in Group #2.

| **Pathway** | **Fold Enrichment** | **P Value** | **Genes** |
| --- | --- | --- | --- |
| TNF signaling pathway | 1.65 | 2E-03 | CEBPB, TNFAIP3, PIK3R3, CXCL1, PIK3CB, PIK3R1, CXCL3, PTGS2, CXCL2, BAG4, CASP8, RPS6KA5, CASP10, CASP3, CCL2, MAPK1, RIPK1, MAP3K8, DNM1L, MAP3K5, JUN, EDN1, MLKL, CHUK, IL15, MMP3, CFLAR, FOS, TNFRSF1B, NFKBIA, ITCH, PIK3CA, IL1B, FAS, TAB3 |
| Cell cycle | 1.50 | 9E-03 | CDKN1C, RB1, HDAC2, CDKN1B, PRKDC, CDC14A, ORC5, CCND3, PTTG1, YWHAQ, CHEK2, MYC, CDC26, RAD21, ORC2, CHEK1, E2F3, BUB3, SKP1, SMAD4, GADD45B, CDC25C, YWHAZ, GADD45G, RBL2, CCNA2, STAG1, WEE1, CDK7, RBL1, DBF4, CCNE2, CDK2, CDK1, MDM2, ANAPC4, MAD2L1 |
| Steroid biosynthesis | 2.52 | 1E-02 | CYP27B1, EBP, SOAT1, SOAT2, CYP51A1, DHCR24, CEL, DHCR7, HSD17B7, LIPA |
| MAPK signaling pathway | 1.29 | 2E-02 | HSPB1, FGF1, RPS6KA6, RPS6KA5, MECOM, MYC, RPS6KA2, CASP3, MAP3K8, MAP3K6, MAP3K4, MAP3K5, DUSP4, PDGFRA, DUSP5, DUSP3, CHUK, DUSP1, IL1R1, PLA2G4B, RRAS2, PLA2G4A, PRKCA, FOS, MAPK8IP2, TGFBR1, DUSP6, TGFBR2, DUSP7, CACNB2, PPM1A, PPM1B, TRAF6, IL1B, RAPGEF2, RAF1, CRK, SOS2, RASGRP2, RASGRP4, CACNA1H, CACNA1G, CDC42, CACNG8, PPP3R1, RAP1A, MAPK7, FLNA, MAPK1, FLNB, MAP4K3, FGF22, NTRK2, JUN, GADD45B, BDNF, NFATC3, BRAF, HSPA2, GADD45G, PPP5C, FGF14, FAS, MAP3K13, FGFR4 |
| mTOR signaling pathway | 1.65 | 3E-02 | PRKAA1, PRKAA2, CAB39, CAB39L, IRS1, RPS6, PIK3R3, BRAF, PRKCA, TSC1, PIK3CB, PIK3R1, MTOR, RPS6KA6, PIK3CA, RPS6KA2, ULK2, MAPK1, RICTOR |
| Phosphatidylinositol signaling system | 1.44 | 4E-02 | DGKG, PI4K2B, MTMR2, DGKB, ITPR1, MTMR8, PIK3R3, PIK3CB, PIK3R1, MTM1, IMPA1, IMPA2, PIP4K2A, PLCE1, PLCG1, IP6K3, IPMK, ITPK1, PRKCA, OCRL, INPP4B, SYNJ1, PIK3CA, IMPAD1, PIK3C3, CALM1, DGKI, DGKH |
| Vascular smooth muscle contraction | 1.38 | 5E-02 | GUCY1B3, RAMP2, ROCK1, ROCK2, NPR1, ITPR1, ADCY3, ADCY2, ADRA1B, ACTG2, PPP1CB, PPP1CC, KCNMB3, MAPK1, PPP1R14A, PTGIR, PLA2G4B, PLA2G4A, PRKCA, BRAF, ACTA2, ADORA2A, ADORA2B, GNAQ, KCNMA1, AGTR1, MRVI1, RAF1, CALM1, PPP1R12B, RAMP1, PPP1R12C |
| VEGF signaling pathway | 1.49 | 8E-02 | PLA2G4B, PXN, NFATC2, PLA2G4A, HSPB1, PIK3R3, PRKCA, PIK3CB, PIK3R1, PTGS2, PTK2, CDC42, PPP3R1, PIK3CA, KDR, MAPK1, PLCG1, RAF1 |
| TGF-beta signaling pathway | 1.38 | 1E-01 | TGIF1, SMAD4, ROCK1, ZFYVE9, NOG, BMP8A, SMAD9, INHBB, GDF5, LTBP1, ACVR2B, DCN, TGFBR1, BMP6, TGFBR2, GDF7, PPP2CA, BMP2, RBL1, PPP2R1B, MYC, MAPK1, SKP1 |
| ECM-receptor interaction | -2.37 | 5E-09 | LAMA5, TNXB, LAMA2, SDC4, LAMA1, ITGA2B, TNC, LAMC2, THBS2, THBS4, THBS3, COMP, VTN, RELN, SV2A, DAG1, TNR, ITGB8, ITGB6, CD36, COL27A1, LAMB3, LAMB2, ITGA3, HSPG2, COL1A1, COL3A1, COL1A2, COL4A2, COL5A1, COL4A1, COL6A2, COL5A2, COL6A1, ITGA11, ITGA8, COL4A5, COL6A3, ITGA7, COL6A6, COL6A5, AGRN, CD44, ITGA9 |
| Axon guidance | -2.14 | 2E-09 | SEMA5A, ROBO3, LRRC4, RND1, RGS3, NCK2, RAC2, RAC3, PLXNC1, EPHB2, HRAS, EPHB1, EPHB4, EPHB3, SEMA6B, EPHA5, UNC5B, EPHA8, UNC5C, UNC5D, RHOD, RASA1, MET, NGEF, EPHA3, EPHA2, SEMA3D, SEMA3B, NTN4, SEMA3E, SEMA3F, EFNA4, NTN3, GNAI2, EFNB2, EFNB1, PAK1, EFNB3, ABL1, PAK6, PLXNA1, SLIT3, FYN, SRGAP3, PAK3, SRGAP2, LRRC4C, NTNG1, NTNG2, SEMA4A, LIMK1, SEMA4F, SEMA4G, L1CAM, EFNA3, CXCL12, PLXNB3, PLXNB2 |
| ABC transporters | -1.81 | 2E-02 | ABCD4, ABCA2, ABCD2, ABCB1, ABCA3, ABCB4, TAP2, TAP1, ABCA7, ABCC9, ABCB9, ABCB11, ABCA12, ABCC11, ABCA13, ABCG1, CFTR |
| Notch signaling pathway | -1.75 | 2E-02 | JAG2, NUMBL, JAG1, CTBP1, PSEN2, DTX1, DTX2, RFNG, DLL1, DLL3, APH1A, DLL4, LFNG, NCSTN, DVL1, DVL2, DVL3, HES5 |
| Cell adhesion molecules (CAMs) | -1.71 | 4E-05 | CNTNAP2, CNTNAP1, ITGB2, ICAM2, LRRC4, ITGAL, PVR, CLDN1, PTPRF, ICAM1, CDH5, CDH3, CDH2, CDH1, MPZ, ITGB8, NEO1, CD34, HLA-B, HLA-C, HLA-A, HLA-F, VCAN, CLDN7, ITGA8, CDH15, ITGA9, NLGN3, SELPLG, NLGN2, SDC4, NRXN1, SDC2, SDC3, NRXN2, GLG1, HLA-DMA, LRRC4C, CD99, JAM2, LRRC4B, CD276, NTNG1, NTNG2, CADM3, VCAM1, CADM1, L1CAM, CD4, OCLN, PTPRC, SIGLEC1 |
| Gap junction | -1.65 | 4E-03 | ADCY4, HTR2B, HTR2A, ADCY1, ADCY8, ADCY7, GRM1, GNAI2, ADCY5, TUBA1C, TUBB6, GRM5, TUBA1A, TUBB3, PDGFC, PRKACA, HRAS, PRKG1, MAP2K5, GUCY1A3, GUCY1A2, MAP2K1, PRKCB, EGF, TUBB, PLCB4, ADCY9, GNAS, GRB2, SOS1, PLCB2 |
| Inflammatory mediator regulation of TRP channels | -1.57 | 6E-03 | CALML6, CAMK2A, ADCY4, HTR2B, PIK3R2, HTR2A, ADCY1, ADCY8, ADCY7, ADCY5, PLCG2, BDKRB2, BDKRB1, PRKACA, MAP2K6, ASIC1, MAP2K3, NTRK1, PRKCH, PRKCB, PRKCE, PLA2G4C, IGF1, NGF, MAPK10, MAPK11, PLCB4, ADCY9, TRPV4, GNAS, PRKCQ, PLCB2, CALM2 |
| Long-term depression | -1.48 | 7E-02 | GUCY1A3, GUCY1A2, GRIA2, GNAZ, MAP2K1, GRID2, PRKCB, PLA2G4C, CACNA1A, IGF1, GRM1, GNAI2, PLCB4, GNA12, GNAS, HRAS, PLCB2, GRIA3, PRKG1 |
| Ras signaling pathway | -1.43 | 1E-03 | CALML6, RASGRF2, ELK1, FGF7, RASSF5, RAC2, RAC3, PRKACA, HRAS, MAP2K1, KSR1, PRKCB, HGF, PLA2G4C, NGF, PLA2G16, TIAM1, MRAS, RASA1, KIT, SOS1, MET, PLA1A, EPHA2, SHC4, SHC2, SHC1, PLA2G1B, PIK3R2, PLD1, RASGRP1, EFNA4, RELA, RASGRP3, PAK1, GNG4, GNG7, PDGFC, ABL1, PLCG2, PAK6, PAK3, RALGDS, ANGPT4, NGFR, ANGPT2, EGF, BAD, VEGFB, VEGFC, IGF1, GRIN2B, GNG11, NFKB1, VEGFA, MAPK10, FGF17, EFNA3, KITLG, FGF18, GRB2, GNB5, TEK, CALM2, FGF12, FGFR3, RGL2, FGFR2, FGFR1 |
| PI3K-Akt signaling pathway | -1.42 | 5E-05 | EPO, ITGA2B, TNC, COMP, STK11, FGF7, CCND2, CCND1, CREB3L4, CREB3L1, TNR, IFNAR2, MAP2K1, COL27A1, IL4R, HGF, F2R, COL4A2, COL4A1, DDIT4, COL4A5, MTCP1, TP53, EPHA2, ATF4, PIK3R2, EFNA4, VTN, PDGFC, EIF4EBP1, PCK2, NGFR, IGF1, GNG11, NFKB1, FGF17, COL1A1, EFNA3, IL6, CDK6, COL1A2, COL5A1, IL7, CDK4, COL5A2, ITGA11, FGF18, GRB2, GNB5, IL7R, FGF12, FGFR3, FGFR2, FGFR1, CDKN1A, LAMC2, CASP9, MLST8, ITGB8, ITGB6, JAK3, HRAS, ITGA3, NGF, CREB3, COL6A2, COL6A1, KIT, ITGA8, COL6A3, ITGA7, COL6A6, COL6A5, SOS1, MET, ITGA9, CREB5, LAMA5, PHLPP1, TNXB, LAMA2, ATF6B, LAMA1, THBS2, RELA, THBS4, THBS3, RELN, GNG4, GNG7, ANGPT4, ANGPT2, LAMB3, BAD, LAMB2, EGF, VEGFB, VEGFC, PPP2R3B, VEGFA, COL3A1, KITLG, TEK, PKN1, PIK3AP1 |
| cGMP-PKG signaling pathway | -1.42 | 8E-03 | ATF6B, CALML6, ADCY4, ATP1A4, ATP2A3, ATP1A3, ATP1A2, ADRB2, ADCY1, ADCY8, ADCY7, SLC8A1, GNAI2, ADCY5, CREB3L4, CREB3L1, KCNMB2, GNA12, ADORA1, CNGA1, BDKRB2, KCNMB4, CACNA1S, PRKG1, GUCY1A3, GUCY1A2, MAP2K1, MEF2C, KCNJ8, BAD, PRKCE, PDE2A, ATP2B3, ATP2B1, ADRA2B, ADRA2A, CREB3, KCNU1, PLCB4, ADCY9, PDE3A, PDE5A, SLC25A5, PLCB2, CALM2, ATF4, SLC25A6, CREB5 |
| Calcium signaling pathway | -1.38 | 9E-03 | CHRM3, RYR2, CALML6, CHRM5, HTR2B, ATP2A3, HTR2A, HTR4, SLC8A1, GRM1, GRM5, CYSLTR2, BDKRB2, BDKRB1, PRKACA, PRKCB, TNNC1, TNNC2, F2R, TACR1, ITPKB, PLCB4, ADCY9, ITPKA, ORAI2, SLC25A5, PLCB2, SLC25A6, PTGER1, PDE1A, CAMK2A, ADCY4, CACNA1A, ADRB2, ADCY1, ADCY8, CACNA1E, ADCY7, GNA14, PHKG1, ERBB3, TBXA2R, PLCG2, CACNA1S, ATP2B3, ATP2B1, P2RX7, P2RX6, P2RX4, GNAS, PLCD3, CALM2, PLCD4 |
| Regulation of actin cytoskeleton | -1.25 | 5E-02 | CYFIP2, CHRM3, ARPC1B, ITGA2B, ITGB2, CHRM5, ARPC5L, ITGAL, FGF7, BDKRB2, RAC2, BDKRB1, ITGB8, RAC3, PIP4K2B, ITGB6, HRAS, APC2, MAP2K1, ITGA3, INSRR, F2R, VAV2, TIAM1, MRAS, ITGA8, PFN4, ARHGEF4, ITGA7, SOS1, ITGA9, PFN2, PIK3R2, IQGAP2, IQGAP3, FGD1, PAK1, GNA12, PDGFC, PIP5K1A, PAK6, NCKAP1L, PAK3, EGF, LIMK1, BAIAP2, SSH3, MYLPF, FGF17, DIAPH3, ITGA11, FGF18, FGF12, FGFR3, FGFR2, FGFR1 |

Note: Group #2: The pathways Enriched from DEGs between ETH and DEX-treated GC- NR HTM cells; Size: Number of genes enriched.

**Supplementary Table 4c.** List of Enriched Pathways in Group #3.

| **Pathway** | **Fold Enrichment** | **P Value** | **Genes** |
| --- | --- | --- | --- |
| Focal adhesion | 1.86 | 1E-04 | FLT1, SHC3, ROCK1, ROCK2, ITGB4, COL11A1, LAMA3, XIAP, PIK3CB, PIK3R1, MYL12A, ACTB, MYL12B, PPP1CB, CCND3, RAP1A, KDR, SPP1, CAPN2, FLNA, MYL10, FLNB, VAV3, ITGA4, CAV2, CAV1, ITGA2, LAMB4, BRAF, PTK2, PIK3CA, ITGA10, COL4A4, COL4A3, ITGA6, ITGA5, PPP1R12B, TLN1, CRK, SOS2 |
| Phosphatidylinositol signaling system | 2.05 | 2E-03 | DGKG, PI4K2B, MTMR2, DGKB, IPMK, ITPR1, MTMR8, PIK3CB, PIK3R1, MTM1, INPP4B, PIK3CA, IMPA1, IMPA2, IMPAD1, PIP4K2A, PLCE1, PIK3C3, IP6K3, DGKI, DGKH |
| MAPK signaling pathway | 1.59 | 2E-03 | HSPB1, FGF1, RASGRP2, CACNG8, PPP3R1, RAP1A, MAPK7, MECOM, MYC, CASP3, RPS6KA2, FLNA, FLNB, MAP3K8, MAP3K6, MAP4K3, FGF22, MAP3K5, DUSP4, NTRK2, DUSP5, GADD45B, CHUK, DUSP1, NFATC3, BRAF, HSPA2, FOS, TGFBR1, DUSP6, TGFBR2, CACNB2, PPM1A, PPP5C, PPM1B, FGF14, TRAF6, RAPGEF2, FAS, FGFR4, CRK, SOS2 |
| Pathways in cancer | 1.39 | 8E-02 | RB1, WNT2B, PIK3CB, FGF1, ETS1, EDNRB, MECOM, MYC, CASP3, JAK1, CHUK, ITGA2, WNT5A, MITF, FOS, TGFBR1, TGFBR2, MSH6, PIK3CA, CCNE2, COL4A4, TRAF6, COL4A3, AGTR1, RARB, ITGA6, CRK, SOS2, ROCK1, ROCK2, LAMA3, GNAI3, LPAR1, WNT8B, XIAP, PIK3R1, CBL, RASGRP2, HIF1A, FOXO1, FGF22, STAT5A, SMAD4, ZBTB16, PTCH1, FZD6, LAMB4, BRAF, PTK2, NFKBIA, BMP2, FGF14, APC, GNAQ, GNB4, MDM2, FAS |
| TNF signaling pathway | 1.79 | 1E-02 | EDN1, CEBPB, CHUK, TNFAIP3, CFLAR, PIK3CB, PIK3R1, FOS, TNFRSF1B, CXCL2, NFKBIA, ITCH, BAG4, PIK3CA, CASP3, FAS, TAB3, MAP3K8, DNM1L, MAP3K5 |
| p53 signaling pathway | 2.00 | 2E-02 | GADD45B, APAF1, SERPINE1, CCND3, SESN3, CCNE2, CHEK2, CASP3, CCNG2, CCNG1, CHEK1, MDM2, FAS, MDM4 |
| Drug metabolism - cytochrome P450 | 1.97 | 2E-02 | MAOA, ADH1B, ADH1A, MGST1, FMO2, FMO3, FMO4, FMO5, ADH5, ADH6, ADH4, CYP2A6, ALDH3B1, AOX1 |
| Adipocytokine signaling pathway | 1.91 | 3E-02 | PRKAA1, PRKAB2, PRKAA2, CHUK, ACSL1, IRS1, PRKAG2, IRS2, PTPN11, TNFRSF1B, NFKBIA, ACSBG1, JAK2, PPARA |
| Regulation of actin cytoskeleton | 1.46 | 3E-02 | ROCK1, ROCK2, ITGB4, PIK3CB, PIK3R1, FGF1, MYL12A, ACTB, MYL12B, PPP1CB, MYL10, PIP4K2A, FGF22, VAV3, ITGA4, ITGA2, BRAF, ARPC5, SSH2, PTK2, ENAH, DIAPH2, FGF14, APC, PIK3CA, ITGA10, ITGA6, ITGA5, FGFR4, PPP1R12B, CRK, SOS2 |
| Thyroid hormone signaling pathway | 1.58 | 5E-02 | NCOA2, MED1, NCOA3, SLC16A10, PIK3CB, PIK3R1, ATP1B1, ESR1, HIF1A, FOXO1, ACTB, MED13L, MED14, NCOR1, PIK3CA, TBC1D4, MYC, MDM2, PLCE1 |
| Cell cycle | 1.54 | 5E-02 | CDKN1C, RB1, SMAD4, GADD45B, PRKDC, RBL2, CCND3, STAG1, WEE1, RBL1, DBF4, CCNE2, CHEK2, MYC, RAD21, ORC2, CHEK1, MDM2, ANAPC4, BUB3 |
| mTOR signaling pathway | 1.81 | 8E-02 | PRKAA1, PRKAA2, CAB39, PIK3CA, IRS1, RPS6KA2, TSC1, BRAF, RICTOR, PIK3CB, PIK3R1 |
| PI3K-Akt signaling pathway | 1.25 | 9E-02 | PRKAA1, FLT1, PRKAA2, IRS1, ITGB4, COL11A1, LAMA3, LPAR1, BRCA1, PIK3CB, PIK3R1, FOXO3, FGF1, CCND3, BCL2L11, MYC, KDR, SPP1, JAK2, IL6R, FGF22, JAK1, MCL1, ANGPT1, ITGA4, CHUK, ITGA2, INSR, LAMB4, TSC1, PRLR, PTK2, RBL2, FGF14, PIK3CA, CCNE2, ITGA10, COL4A4, COL4A3, GNB4, MDM2, ITGA6, ITGA5, FGFR4, SOS2 |
| TGF-beta signaling pathway | 1.59 | 9E-02 | TGIF1, SMAD4, ROCK1, ZFYVE9, NOG, BMP8A, INHBB, LTBP1, ACVR2B, TGFBR1, TGFBR2, BMP2, RBL1, MYC |
| Axon guidance | -2.25 | 5E-07 | ROBO3, SEMA3D, SEMA3B, SEMA3E, SEMA3F, EFNA4, RND1, NTN3, EFNB1, PAK1, EFNB3, RGS3, ABL1, NCK2, RAC2, PAK6, PLXNA1, RAC3, SLIT3, PLXNC1, SRGAP3, EPHB2, PAK3, LRRC4C, EPHB1, EPHB4, EPHB3, SEMA6B, NTNG2, SEMA4A, UNC5B, SEMA4F, UNC5D, SEMA4G, RHOD, EFNA3, CXCL12, PLXNB2, EPHA3, NGEF, EPHA2 |
| Cell adhesion molecules (CAMs) | -2.01 | 1E-05 | NLGN3, CNTNAP2, CNTNAP1, SELPLG, NLGN2, SDC4, NRXN1, SDC2, SDC3, ITGB2, ICAM2, NRXN2, ITGAL, CLDN1, PTPRF, GLG1, ICAM1, CDH5, HLA-DMA, CDH3, CDH1, LRRC4C, NEO1, CD34, CD99, LRRC4B, CD276, NTNG2, VCAM1, CADM1, HLA-B, HLA-C, HLA-A, HLA-F, CD4, OCLN, PTPRC, CLDN7, SIGLEC1, CDH15, ITGA9 |
| ECM-receptor interaction | -2.24 | 5E-05 | LAMA5, TNXB, SDC4, LAMA1, ITGA2B, TNC, LAMC2, THBS2, THBS4, THBS3, VTN, RELN, SV2A, DAG1, ITGB6, CD36, LAMB3, COL1A1, COL3A1, COL1A2, COL4A2, COL5A1, ITGA11, COL6A3, ITGA7, AGRN, CD44, ITGA9 |
| Ras signaling pathway | -1.60 | 5E-04 | CALML6, RASGRF2, FGF7, RASSF5, RAC2, RAC3, PRKACA, KSR1, PRKCB, HGF, PLA2G4C, NGF, PLA2G16, TIAM1, MRAS, KIT, PLA1A, EPHA2, SHC2, SHC1, PLA2G1B, PIK3R2, EFNA4, PAK1, GNG4, GNG7, PDGFC, ABL1, PLCG2, PAK6, PAK3, RALGDS, ANGPT4, NGFR, ANGPT2, EGF, BAD, VEGFB, VEGFC, IGF1, GRIN2B, GNG11, VEGFA, FGF17, EFNA3, FGF18, GNB5, FGF12, FGFR3, RGL2, FGFR2, FGFR1 |
| Aldosterone synthesis and secretion | -1.98 | 2E-03 | SCARB1, DAGLA, CAMK1D, CALML6, PRKCB, PRKCE, CAMK2A, PDE2A, ADCY4, ADCY1, ADCY7, ADCY5, CREB3, ADCY9, CREB3L4, CYP11A1, CREB3L1, CAMK1, PRKACA, PLCB2, CAMK1G, CREB5, ATF4 |
| Thyroid hormone synthesis | -1.99 | 4E-03 | GPX1, HSPA5, PRKCB, ADCY4, GPX7, ATP1A3, ATP1A2, ADCY1, ADCY7, TSHR, PDIA4, ADCY5, CREB3, ADCY9, CREB3L4, CREB3L1, PRKACA, PLCB2, CREB5, ATF4 |
| Inflammatory mediator regulation of TRP channels | -1.70 | 1E-02 | NTRK1, PRKCH, CALML6, PRKCB, PRKCE, CAMK2A, PLA2G4C, ADCY4, HTR2B, PIK3R2, ADCY1, IGF1, NGF, ADCY7, ADCY5, MAPK11, ADCY9, TRPV4, BDKRB2, PLCG2, BDKRB1, PRKACA, PLCB2, ASIC1 |
| Notch signaling pathway | -2.03 | 2E-02 | JAG2, CTBP1, PSEN2, DTX1, DTX2, RFNG, DLL1, DLL3, APH1A, DLL4, LFNG, DVL1, DVL2, DVL3 |
| Glycine, serine and threonine metabolism | -2.14 | 2E-02 | GAMT, AOC3, AOC2, PSAT1, SHMT2, CHDH, PGAM1, SARDH, PHGDH, SDSL, ALDH7A1, PSPH |
| Calcium signaling pathway | -1.44 | 2E-02 | CHRM3, RYR2, PTGER1, CALML6, PDE1A, CAMK2A, ADCY4, HTR2B, ATP2A3, CACNA1A, ADCY1, CACNA1E, ADCY7, GNA14, CYSLTR2, PHKG1, ERBB3, TBXA2R, PLCG2, BDKRB2, BDKRB1, PRKACA, PRKCB, TNNC1, ATP2B3, TACR1, P2RX7, ITPKB, P2RX6, P2RX4, ADCY9, ORAI2, PLCD3, SLC25A5, PLCD4, PLCB2, SLC25A6 |
| Protein processing in endoplasmic reticulum | -1.44 | 2E-02 | RPN2, PRKCSH, RPN1, RRBP1, HERPUD1, SEC61A1, GANAB, OS9, LMAN2, MAN1C1, FBXO6, BAK1, SIL1, UBQLN4, TXNDC5, PDIA3, SSR4, HSPA5, WFS1, SSR2, FBXO2, HSPA6, UBE2E2, TRAF2, CKAP4, DDOST, PDIA4, DNAJC5, DDIT3, ERP29, HYOU1, P4HB, CALR, MAN1B1, ATF4 |
| Wnt signaling pathway | -1.46 | 3E-02 | CTBP1, TCF7, LEF1, CAMK2A, LRP5, FZD10, FRAT2, WNT11, CCND2, DVL1, DVL2, RAC2, DVL3, RAC3, PRKACA, WNT2, WNT3, WNT4, WNT10B, FZD2, PRKCB, FZD7, CTNNBIP1, FZD8, WNT9A, SFRP4, VANGL2, PLCB2, TP53 |
| ABC transporters | -1.74 | 9E-02 | ABCA2, ABCB1, ABCA3, ABCB4, TAP2, TAP1, ABCA7, ABCB9, ABCC11, ABCG1, CFTR |

Note: Group #3: The pathways Enriched from the overlapping DEGs between Group #1 and Group #2; Size: Number of genes enriched.

**Supplementary Table 4d.** List of Enriched Pathways in Group #4.

| **Pathway** | **Fold Enrichment** | **P Value** | **Genes** |
| --- | --- | --- | --- |
| Axon guidance | 1.94 | 1E-02 | ITGB1, GSK3B, SEMA7A, ARHGEF12, SEMA3C, LRRC4, NTN4, UNC5C, NRAS, RASA1, DPYSL2, CFL2, KRAS, PAK2, MET, SRGAP1, NCK1 |
| Regulation of actin cytoskeleton | 1.59 | 3E-02 | ITGB1, CHRM2, NCKAP1, ARHGEF12, PPP1R12A, RDX, ITGA1, PDGFB, FN1, PIK3CD, ACTN4, IQGAP1, FGF2, FGF5, GNA13, PIKFYVE, NRAS, CFL2, ITGAV, KRAS, WASF1, PAK2, ARHGEF6 |
| ECM-receptor interaction | 2.00 | 3E-02 | ITGB1, LAMA2, COL5A3, ITGA1, CHAD, FN1, TNR, SDC1, COL6A6, ITGAV, GP1BA, COL6A5 |
| Focal adhesion | 1.55 | 5E-02 | ITGB1, GSK3B, PPP1R12A, LAMA2, ITGA1, PDGFB, FN1, PIK3CD, ACTN4, ARHGAP5, IGF1R, MAPK8, COL5A3, CHAD, AKT3, TNR, COL6A6, ITGAV, COL6A5, PAK2, MET, BIRC2 |
| PI3K-Akt signaling pathway | 1.39 | 5E-02 | ITGB1, CHRM2, PHLPP2, ATF2, GSK3B, LAMA2, PDGFB, PIK3CD, FGF2, IGF1R, GHR, FGF5, NRAS, PPP2R3C, CHAD, AKT3, TNR, ITGAV, ITGA1, FN1, PPP2R3A, RPS6KB1, COL5A3, DDIT4, MTCP1, KRAS, PKN2, COL6A6, SGK3, COL6A5, MET, TLR4, TLR2 |
| Adherens junction | 2.04 | 5E-02 | SMAD2, TJP1, YES1, ACTN4, CTNNA2, IQGAP1, WASF1, MAP3K7, MET, IGF1R |
| Protein processing in endoplasmic reticulum | 1.54 | 7E-02 | SEC23A, SEC24A, SAR1B, SEL1L, CUL1, EIF2AK2, YOD1, EIF2S1, UBE2J1, DNAJC3, MAPK8, HSPH1, MAN1A2, MAN1A1, UGGT2, SEC62, ATF6, MBTPS2 |
| T cell receptor signaling pathway | 1.74 | 8E-02 | GSK3B, DLG1, NRAS, AKT3, PIK3CD, KRAS, MAP3K7, PAK2, RASGRP1, LAT, MALT1, NCK1 |
| Cell cycle | 1.64 | 8E-02 | SMAD2, GSK3B, CCNH, CUL1, SMC3, CDC14B, CCNB3, CCNA1, STAG2, ORC3, CDC27, ATM, BUB1, ATR |
| Calcium signaling pathway | -1.98 | 1E-05 | CAMK2B, RYR1, MYLK2, CHRM1, PDE1B, CHRNA7, PTAFR, CACNA1B, CACNA1D, ADRA1D, ADRB1, ATP2A1, CALML3, ADCY2, GRPR, ADRA1A, SLC8A2, CACNA1G, MYLK4, PPP3CA, HRH1, GNA15, CYSLTR1, PLN, PHKG2, ERBB2, CD38, DRD1, NOS1, NTSR1, PDGFRA, NOS3, TACR2, AVPR1A, GRIN2C, GRIN1, LTB4R2, ADORA2B, STIM2, P2RX2, P2RX1, CAMK4, CALM3 |
| Type I diabetes mellitus | -3.33 | 1E-05 | CD86, HLA-DRB5, PTPRN2, PTPRN, GAD1, PRF1, HLA-DMB, IL1B, HLA-DPB1, HLA-DRA, IL12A, HLA-DOA, HLA-DQA2, HLA-DQA1, HLA-DRB1, HLA-DPA1, HLA-DQB1 |
| Cell adhesion molecules (CAMs) | -1.85 | 7E-04 | CD86, ITGAM, F11R, SPN, HLA-DMB, ITGB7, HLA-DOA, HLA-DQA2, ICOSLG, HLA-DQA1, HLA-DPA1, HLA-DRB5, SELE, SELP, CLDN11, CD2, CLDN6, CLDN5, CLDN4, CLDN3, SELL, CD6, CD8A, CLDN8, HLA-DPB1, PECAM1, HLA-DRA, CNTN2, ESAM, CD22, HLA-DRB1, HLA-DQB1 |
| B cell receptor signaling pathway | -2.03 | 7E-03 | LYN, SYK, NFATC2, PIK3R3, NFATC1, PIK3CG, VAV1, PIK3R5, CD79B, CD79A, PPP3CA, AKT2, BLNK, PTPN6, FCGR2B, CD22, CARD11 |
| TNF signaling pathway | -1.69 | 2E-02 | MLKL, RIPK3, IL15, MMP3, LIF, PIK3R3, CXCL1, CXCL3, SELE, MMP9, PIK3CG, MAPK12, PIK3R5, TNFRSF1A, CXCL10, SOCS3, CASP10, IL1B, AKT2, CCL5, CCL2, PGAM5 |
| Chemokine signaling pathway | -1.37 | 6E-02 | CCL14, CCL13, CXCL6, GSK3A, CXCL9, NCF1, WAS, CXCR4, PIK3R3, CXCL1, ADCY2, ARRB2, CXCL13, CXCL3, CXCL14, PIK3CG, PIK3R5, PREX1, AKT2, CCL5, CCL2, CCL19, CCR1, LYN, PRKCD, VAV1, FGR, CXCL10, HCK, CXCL11, CCL26 |

Note: Group #4: Uniquely expressed GC-R HTM cells specific pathways enriched from DEGs of Group #3 minus Group #1; Size: Number of genes enriched.

**Supplementary Table 4e.** List of Enriched Pathways in Group #5.

| **Pathway** | **Fold Enrichment** | **P Value** | **Genes** |
| --- | --- | --- | --- |
| Long-term depression | 2.31 | 9E-03 | RYR1, GUCY1B3, PLA2G4B, PLA2G4A, PRKCA, GNAI1, PPP2CA, GNAO1, PPP2R1B, PRKG2, MAPK1, NOS1, RAF1 |
| Vascular smooth muscle contraction | 1.82 | 1E-02 | GUCY1B3, PTGIR, RAMP2, NPR1, PLA2G4B, PLA2G4A, ADCY3, PRKCA, ADCY2, ACTG2, PPP1CC, ADORA2A, ADORA2B, KCNMA1, MAPK1, MRVI1, CALM1, RAF1, PPP1R12C, RAMP1 |
| Inflammatory mediator regulation of TRP channels | 1.85 | 2E-02 | PTGER4, IL1R1, PTGER2, PLA2G4B, PLA2G4A, ADCY3, TRPV3, PIK3R3, PRKCA, ADCY2, ALOX12, PPP1CC, IL1B, F2RL1, PLCG1, CALM1, ASIC3 |
| Fatty acid metabolism | 2.22 | 3E-02 | CPT1A, FASN, OXSM, ACSL6, ACSL5, ELOVL6, HADH, CPT1B, ACAT2, ACAT1 |
| VEGF signaling pathway | 1.92 | 6E-02 | CDC42, PLA2G4B, PXN, NFATC2, PIK3R3, PLA2G4A, MAPK1, PRKCA, PLCG1, RAF1, PTGS2 |
| Oxytocin signaling pathway | 1.49 | 7E-02 | RYR1, GUCY1B3, JUN, NPR1, PLA2G4B, KCNJ14, NFATC2, PLA2G4A, ADCY3, PRKCA, ADCY2, PTGS2, GNAI1, GNAO1, PPP1CC, RGS2, CAMK4, MAPK1, CALM1, RAF1, PPP1R12C |
| Calcium signaling pathway | 1.43 | 7E-02 | RYR1, PDGFRA, PTGFR, PDE1C, ATP2A2, ADCY3, ATP2A1, PRKCA, ADCY2, GRIN2C, CACNA1H, SLC8A2, CACNA1G, PLN, GNAL, STIM1, ADORA2A, ADORA2B, CAMK4, P2RX1, VDAC3, NOS1, PLCG1, CALM1 |
| ECM-receptor interaction | -2.62 | 9E-04 | COL27A1, LAMA2, LAMB2, ITGA3, HSPG2, COMP, COL4A1, COL6A2, COL5A2, COL6A1, ITGA8, COL4A5, TNR, ITGB8, COL6A6, COL6A5 |
| Focal adhesion | -1.94 | 1E-03 | SHC4, LAMA2, ELK1, COMP, CCND1, TNR, ITGB8, FYN, HRAS, MAP2K1, COL27A1, LAMB2, ITGA3, PARVA, MYLPF, MAPK10, COL4A1, COL6A2, COL5A2, COL6A1, ITGA8, COL4A5, GRB2, COL6A6, TLN2, COL6A5, SOS1, MET |
| Gap junction | -2.43 | 3E-03 | MAP2K1, HTR2A, ADCY8, GRM1, GNAI2, GRM5, TUBB6, TUBA1A, PLCB4, GNAS, GRB2, SOS1, HRAS, MAP2K5, PRKG1 |
| Axon guidance | -1.91 | 2E-02 | NTNG1, SEMA5A, EPHA5, EPHA8, LIMK1, LRRC4, NTN4, UNC5C, L1CAM, GNAI2, EFNB2, RASA1, PLXNB3, FYN, SRGAP2, HRAS, MET |
| PI3K-Akt signaling pathway | -1.45 | 2E-02 | PHLPP1, LAMA2, ATF6B, EPO, RELA, COMP, CCND1, EIF4EBP1, TNR, ITGB8, HRAS, JAK3, MAP2K1, COL27A1, LAMB2, ITGA3, F2R, NFKB1, KITLG, CDK6, COL4A1, COL6A2, COL5A2, COL6A1, DDIT4, ITGA8, COL4A5, MTCP1, GRB2, COL6A6, COL6A5, PKN1, TEK, SOS1, MET |
| MAPK signaling pathway | -1.52 | 3E-02 | PPP5D1, ARRB1, ECSIT, RASGRP1, ELK1, RELA, RASGRP3, DUSP10, GNA12, NTF3, CACNA1S, CD14, HRAS, MAP2K5, MAP2K6, MAP2K3, TGFB2, MAP2K1, MEF2C, DUSP2, MAP3K1, NFKB1, MAPK10, RASA1, GRB2, SOS1, MAP3K14 |
| Pathways in cancer | -1.38 | 4E-02 | LAMA2, EPAS1, BRCA2, ADCY8, RASGRP1, RELA, GNAI2, RASGRP3, CCND1, TPR, GNA12, CTNNA1, VHL, CTNNA2, SKP2, HRAS, APC2, TCF7L2, TGFB2, MAP2K1, TCF7L1, CDKN2A, LAMB2, ITGA3, F2R, STAT3, NFKB1, PML, MAPK10, KITLG, PLCB4, CDK6, COL4A1, GNAS, COL4A5, GRB2, SOS1, MET |
| Insulin secretion | -1.85 | 7E-02 | KCNU1, PLCB4, ATF6B, KCNMB2, ATP1A4, GNAS, KCNMB4, KCNN2, CACNA1S, ADCY8, TRPM4 |
| Cell cycle | -1.61 | 9E-02 | TGFB2, CDKN2A, BUB1B, TTK, CDC7, SMC1B, CCNA1, CDK6, CCND1, ATM, SKP2, E2F5, BUB1, MAD1L1 |

Note: Group #5: Uniquely expressed GC-NR HTM cells specific pathways enriched from DEGs of Group #3 minus Group #2; Size: Number of genes enriched.

**Supplementary Table 5**: Comparison of Differentially Expressed Genes in TM Cells Treated with Glucocorticoid from Previous Reports and the Present Study

| **Study Design** | **Use of Postmortem Tissue** | **Age of the donors (Yrs)** | **Number of**  **Donors used** | **Passage of Cells** | **Dexamethasone**  **Treatment(nM)** | **Duration of Exposure (d)** | **Control medium** | **History of**  **glaucoma** | **Microarray/RNA Seq** | **List of Up-regulated genes** | **References** |
| --- | --- | --- | --- | --- | --- | --- | --- | --- | --- | --- | --- |
| Human Cultured TM cells | yes | 7,16,21 and 28 | 4 | 4-5 | 100 | 7 | 0.1% Ethanol | Not specified | cDNA | RGC32, OCA2, ANGPTL7, MYOC, FKBP5, SAA1, ZBTB16 | #8 |
| Human Cultured TM cells | Yes | unknown | 2 | 4-6 | 100 | 10 | 0.1% Ethanol | No history of glaucoma | Affymetrix Human Genome U95Av2 gene Chip microarrays | SERPINA3, AKR1C1, IGFBP2, MYOC, PEDF, PCOLCE, AKR1C3, PTGDS, CHI3L1, ACTG2, CDT6, ASS, CTP, APOD, AP1G1 | #9 |
| Human Cultured TM cells |  | unknown |  | 8 | 100 | 10 | Water in DMEM medium | Not specified | MicroMax Human cDNA System I | GAS1, CDH4, MT1L, CST3, TSC22, LDHA, IGFBp2, TAGLN | #10 |
| Human Cultured TM cells | Yes | 12,16 and 17 | 3 | 5 | 100 | 21 | DMEM medium containing 10% fetal calf serum | No history of glaucoma | Affymetrix Human Genome U133A array | MYOC, DCN, IGFBP2 | #11 |
| Human Cultured TM cells | Yes | 52 | 1 | 8 | 100 | 7 | 0.0025% and 0.025% Benzyl alcohol | No history of glaucoma | Standford Human cDNA SHEW | GAS1, CDH4, MT1L, CST3, ATF4, ASNS/TS11, CHOP, HSPA5 | #4 |
| Human Cultured TM cells | Yes | 3 months old and 35 | 2 | 3-6 | 1000 | 24h | 0.1% DMSO | Not specified | Agilent-014850 whole Human Genome Microarray 4X44K G4112F | RGC32, OCA2, SAA1, ANGPTL7, ITGA10, LSP1, KCNB1, FKBP5, KLF15, ANGPTL4, MYOC, ZBTB16, PF4 | #5 |
| Human Cultured TM cells | Yes | Not specified | 3 | Not specified | 100 | 14 | DMEM with 10% fetal calf serum | Not specified | Agilent -028004 SurePrint G3 Human  Gene Expression 8X60 K Microarray 039381 | FKBP5, ZBTB16, SCNN1A, SAA1 | #6 |
| Human Cultured TM cells | Yes | 27 | 1 | 2 | 500 | 6 | 0.1% ethanol | Not specified | Affymetrix Human Gene 1.0 ST Array | ITGB3, CHN1, PIK3R1, MFGE8 | #7 |
| Bovine – Perfusion organ culture system  -BTM cells with known GC responsiveness used | Yes-Bovine donor eyes | Adult cows | 6 donors (paired eyes) | >3 | 100 | 7 | 0.1% ethanol | Not specified | Illumina Human Methylation 450K Chips | DKK1, HMGA2, SOD2, STEAP4, SLC7A8 | #13 |
| Human – Perfusion organ culture system  -HTM cells with known GC responsiveness | Yes | 67 (mean) | 8 donors | 3-4 | 100 | 7 | 0.1% Ethanol | No history of glaucoma | Illumina Next Seq 500 | ZBTB16, SAA1, MYOC, OCA2, H19, APOD, SAA2, HIF3A, PRODH, FKBP5, LSP1, ANGPTL7, CPM. STEAP4, MT2A, IRF7 | Present Study |

**Supplementary Table 6a**. Comparison of Pathways Involved in DEX treated TM Cells from Previous Reports and the Present Study

| **Effect of DEX on TM cells** | | **Present Study** |
| --- | --- | --- |
| **Human Studies Combined*** | **Bovine**^ɫ^ |  |
| Collagen | Collagen | - |
| ECM | ECM | - |
| Adhesion | Adhesion | Adhesion |
| WNT signaling | WNT signaling | WNT signaling |
| Inflammation | Inflammation | - |
| Adipogenesis | Adipogenesis | - |
| Glucose metabolism | Glucose metabolism | - |
| NF-κB | NF-κB | - |
| Apoptosis | Apoptosis | - |
| GPCR | GPCR | - |
| Oxidative stress | Oxidative stress | - |
|  |  | Phosphatidylinositol signaling system |
|  |  | TGF Beta Signaling Pathway |
|  |  | P53 Signaling Pathway |
|  |  | mTOR signaling pathway |
|  |  | Cell cycle |
|  |  | Adherens junction |
|  |  | Regulation of Actin Cytoskeleton |
|  |  | TNF signaling pathway |
|  |  | MAPK Signaling Pathway |
|  |  | Pathways in Cancer |
|  |  | PI3K-Akt signaling pathway |
|  |  | Cell Adhesion Molecules Cams |
|  |  | Calcium signaling pathway |
|  |  | Axon Guidance |
|  |  | Inflammatory mediator regulation of TRP channels |
|  |  | ECM-receptor interaction |
|  |  | cAMP signaling pathway |
|  |  | Ras signaling pathway |
|  |  | Steroid biosynthesis |
|  |  | Vascular smooth muscle contraction |
|  |  | VEGF Signaling Pathway |
|  |  | ABC transporters |
|  |  | Notch signaling pathway |
|  |  | Gap junction |
|  |  | cGMP-PKG signaling pathway |

***Note****:* **^*^***Ref. No.*4-11, ^ϯ^ Ref.No.13, 23.

**Supplementary Table 6b**. Comparison of Pathways Enriched in Responder and Non-Responder HTM Cells from Previous Report and the Present Study.

| **Responder vs. non-responders Bovine** ^‡^ | **Present Study** | |
| --- | --- | --- |
|  | **Responder** | **Non-Responder** |
| Collagen | - |  |
| ECM | - |  |
| Adhesion | Adhesion |  |
| WNT signaling | WNT signaling |  |
| Inflammation | - |  |
| Adipogenesis | - |  |
| Glucose metabolism | - |  |
| NF-κB | - |  |
| Apoptosis | - |  |
| Cell cycle | Cell cycle |  |
| Senescence | - |  |
| Oxidative stress | - |  |
|  | Axon guidance | Long-term depression |
|  | Regulation of actin cytoskeleton | Vascular smooth muscle contraction |
|  | ECM receptor interaction | Inflammatory mediator regulation of TRP channels |
|  | Focal adhesion | Fatty acid metabolism |
|  | PI3K-Akt signaling pathway | VEGF signaling pathway |
|  | Adherens junction | Oxytocin signaling pathway |
|  | Protein processing in endoplasmic reticulum | Calcium signaling pathway |
|  | T cell receptor signaling pathway | ECM-receptor interaction |
|  | Calcium signaling pathway | Focal adhesion |
|  | Cell adhesion molecules (CAMs) | Gap junction |
|  | B cell receptor signaling pathway | Axon guidance |
|  | TNF signaling pathway | PI3K-Akt signaling pathway |
|  | Chemokine signaling pathway | MAPK signaling pathway |
|  |  | Pathways in cancer |
|  |  | Insulin secretion |
|  |  | Cell cycle |

***Note****:* ^‡^ *Ref. No.*13, 23

**Supplementary Table 7:** Characteristics of Human Donor Eyes Used for the Present Study

| **DEX Treatment (nM)** | **Code** | **Age** | **Sex** | **Cause of Death** | **Time B/W Death & Enucleation (h)** | **Time B/W Enucleation & Culture (h)** | **Experiment**  **Eye** | **HOCAS/ Culture** | **Treatment** | **Remarks** |
| --- | --- | --- | --- | --- | --- | --- | --- | --- | --- | --- |
| 100nM (N=16) | OCHD18-29 | 76 | F | Natural | 0.45 | 30.5 | OD | HOCAS | DEX | Data included |
|  |  |  |  |  |  |  | OS | Cell culture | ETH/DEX | TM Culture established* |
|  | OCHD18-49 | 48 | M | Heart Disease | 2.5 | 30.66 | OD | HOCAS | DEX | Data included |
|  |  |  |  |  |  |  | OS | Cell culture | ETH/DEX | TM Culture established |
|  | OCHD18-52 | 55 | F | Vascular accident | 2.75 | 67.91 | OD | HOCAS | DEX | Data included |
|  |  |  |  |  |  |  | OS | Cell culture | ETH/DEX | TM Culture established* |
|  | OCHD18-53 | 67 | M | Diabetes | 4.33 | 27 | OD | HOCAS | DEX | Data included |
|  |  |  |  |  |  |  | OS | Cell culture | ETH/DEX | TM Culture established* |
|  | OCHD18-56 | 82 | M | Cardiac arrest | 4 | 23 | OD | HOCAS | DEX | Data included |
|  |  |  |  |  |  |  | OS | Cell culture | ETH/DEX | TM Culture established* |
|  | OCHD18-57 | 80 | F | Respiratory disease | 3 | 28.5 | OD | HOCAS | DEX | Data included |
|  |  |  |  |  |  |  | OS | Cell culture | ETH/DEX | TM Culture established |
|  | OCHD19-01 | 73 | M | Natural | 4.5 | 30.8 | OD | HOCAS | DEX | Data included |
|  |  |  |  |  |  |  | OS | Cell culture | ETH/DEX | TM Culture established |
|  | OCHD19-02 | 66 | M | Road traffic injury | 2.75 | 8.33 | OD | HOCAS | DEX | Data included |
|  |  |  |  |  |  |  | OS | Cell culture | ETH/DEX | TM Culture established* |
|  | OCHD19-03 | 65 | F | Heart Disease | 5.8 | 47.33 | OD | HOCAS | DEX | Data included |
|  |  |  |  |  |  |  | OS | Cell culture | ETH/DEX | TM Culture established* |
|  | OCHD19-04 | 72 | M | Heart Disease | 3.5 | 41 | OD | HOCAS | DEX | Data included |
|  |  |  |  |  |  |  | OS | Cell culture | ETH/DEX | TM Culture established* |
|  | OCHD20-04 | 65 | M | Chronic obstructive pulmonary disease | 0.33 | 28.5 | OD | HOCAS | DEX | Data included |
|  |  |  |  |  |  |  | OS | Cell culture | ETH/DEX | TM Culture established |
|  | OCHD20-05 | 50 | M | Kidney failure | 2 | 71.58 | OD | HOCAS | DEX | Data included |
|  |  |  |  |  |  |  | OS | Cell culture | ETH/DEX | TM Culture established |
|  | OCHD20-06 | 59 | M | Chronic Kidney Disease | 0.5 | 29.16 | OD | HOCAS | DEX | Data included |
|  |  |  |  |  |  |  | OS | Cell culture | ETH/DEX | TM Culture established |
|  | OCHD20-09 | 46 | M | Accidental fall | 2.25 | 31 | OD | HOCAS | DEX | Data included |
|  |  |  |  |  |  |  | OS | Cell culture | ETH/DEX | TM Culture established |
|  | OCHD20-10 | 38 | M | Chronic Kidney Disease | 1.41 | 47.5 | OD | HOCAS | DEX | Data included |
|  |  |  |  |  |  |  | OS | Cell culture | ETH/DEX | TM Culture established |
|  | OCHD20-14 | 56 | M | Heart Disease | 3.91 | 72.25 | OD | HOCAS | DEX | Data included |
|  |  |  |  |  |  |  | OS | Cell culture | ETH/DEX | TM Culture established |
|  | **HTM Cell used for RNA-Sequencing** | | | | | | | | | |
|  | OCHD18-39 | 82 | F | Respiratory disease | 0.75 | - | OS | Cell culture | ETH/DEX | TM Culture established* |

A total of 16 paired eyes were used for the study with the mean (±) SD age was 62.38 ± 12.82 years. The mean (±) SD elapsed time between (b/w) death and enucleation was 2.75 ± 1.58 h and the mean elapsed time between enucleation and culture was 38.44 ± 18.39 h. DEX- Eyes treated with dexamethasone; ETH/DEX- HTM Cells treated with either 0.1% ethanol or dexamethasone. * HTM Cells used for RNA-sequencing.

**Supplementary Table 8**. List of Genes Selected from RNA Sequencing for Validation by RT^2^- PCR Array

| **S.No** | **Ref seq** | **Symbol** | **Description** |
| --- | --- | --- | --- |
| 1 | NM_000681 | ADRA2A | Adrenergic, alpha-2A-, receptor |
| 2 | NM_004673 | ANGPTL1 | Angiopoietin-like 1 |
| 3 | NM_001647 | APOD | Apolipoprotein D |
| 4 | NM_031911 | C1QTNF7 | C1q and tumor necrosis factor related protein 7 |
| 5 | NM_004355 | CD74 | CD74 molecule, major histocompatibility complex, class II invariant chain |
| 6 | NM_001276 | CHI3L1 | Chitinase 3-like 1 (cartilage glycoprotein-39) |
| 7 | NM_022131 | CLSTN2 | Calsyntenin 2 |
| 8 | NM_001855 | COL15A1 | Collagen, type XV, alpha 1 |
| 9 | NM_001874 | CPM | Carboxypeptidase M |
| 10 | NM_004419 | DUSP5 | Dual specificity phosphatase 5 |
| 11 | NM_006350 | FST | Follistatin |
| 12 | NM_000618 | IGF1 | Insulin-like growth factor 1 (somatomedin C) |
| 13 | NM_000612 | IGF2 | Insulin-like growth factor 2 (somatomedin A) |
| 14 | NM_000261 | MYOC | Myocilin, trabecular meshwork inducible glucocorticoid response |
| 15 | NM_015039 | NMNAT2 | Nicotinamide nucleotide adenylyltransferase 2 |
| 16 | NM_000905 | NPY | Neuropeptide Y |
| 17 | NM_005010 | NRCAM | Neuronal cell adhesion molecule |
| 18 | NM_006180 | NTRK2 | Neurotrophic tyrosine kinase, receptor, type 2 |
| 19 | NM_000275 | OCA2 | Oculocutaneous albinism II |
| 20 | NM_002612 | PDK4 | Pyruvate dehydrogenase kinase, isozyme 4 |
| 21 | NM_002820 | PTHLH | Parathyroid hormone-like hormone |
| 22 | NM_004103 | PTK2B | PTK2B protein tyrosine kinase 2 beta |
| 23 | NM_014059 | RGCC | Regulator Of Cell Cycle |
| 24 | NM_000331 | SAA1 | Serum amyloid A1 |
| 25 | NM_006922 | SCN3A | Sodium channel, voltage-gated, type III, alpha subunit |
| 26 | NM_016932 | SIX2 | SIX homeobox 2 |
| 27 | NM_012391 | SPDEF | SAM pointed domain containing ets transcription factor |
| 28 | NM_024636 | STEAP4 | STEAP family member 4 |
| 29 | NM_003256 | TIMP4 | TIMP metallopeptidase inhibitor 4 |
| 30 | NM_170744 | UNC5B | Unc-5 homolog B |
| 31 | NM_004385 | VCAN | Versican |
| 32 | NM_006006 | ZBTB16 | Zinc finger and BTB domain containing 16 |
